# CADD-SV v2.0: An Extended Framework for Structural Variant Deleteriousness Prediction Using Coordinate- and Sequence-Based Features

**DOI:** 10.64898/2026.07.08.736040

**Authors:** Orazio Catona, Martin Kircher

## Abstract

Structural variants are a major source of genomic variation and contribute to human disease and evolution through diverse mechanisms, yet their functional interpretation remains challenging. We present CADD-SV v2.0, an improved machine learning framework for scoring SV deleteriousness that expands on the original CADD-SV implementation. This version introduces a unified Random Forest model trained on an expanded set of proxy-neutral and proxy-deleterious variants drawn from human and non-human primate genomes. The model integrates updated genomic annotations, including constraint metrics, regulatory elements, and chromatin architecture features. It scores Deletions, Insertions, Duplications and Inversions based on a single scoring framework that uses both the variant and its flanking regions. To complement this framework, we also explore sequence-based annotations derived from SegmentNT, a deep learning model that provides functional predictions from DNA sequence at nucleotide resolution. Our analysis evaluated whether sequence-derived functional signals can provide additional information for SV prioritization and whether additional models with these features alone or in combination with previous coordinate-based annotations can be used. CADD-SV v2.0 outperforms its previous version and other tools in prioritizing deleterious variants across major SV types, including some previously unsupported, and substantially improves the computational workflow, increasing predictive power for genome-wide SV interpretation.

## Introduction

Structural variants (SVs), typically defined as genomic alterations spanning 50 base pairs (bp) or more, represent a major and diverse class of genetic variation in the human genome. In contrast to single-nucleotide variants (SNVs), SVs vary widely in type, length, and genomic complexity. They can involve simple events such as deletions and insertions, but also encompass duplications, inversions, translocations, and even more complex rearrangements that are a combination of those. These events collectively account for more altered base pairs per genome than SNVs and have the capacity to profoundly reshape genomic structure and function (1). SVs can influence DNA at multiple levels. They may disrupt coding sequences, alter gene dosage, affect regulatory elements, or reposition chromatin domains. Importantly, their impact is not limited to genic regions: SVs can interfere with enhancers, silencers, and insulators, alter three-dimensional chromatin interactions, and modify epigenomic states. Consequently, SVs have been implicated in a wide range of phenotypic outcomes and disease mechanisms. They are known to play a role in Mendelian disorders, contribute to complex traits, and are frequently observed as driver events in cancers. At the same time, SVs also shape population diversity and have played a critical role in primate evolution, influencing gene family expansions, genomic innovation, and adaptation (2).

In recent years, significant advances in sequencing technologies, particularly the advent and refinement of long-read sequencing methods, have substantially improved our ability to detect and characterize SVs in the human genome. Unlike traditional short-read sequencing, long-read platforms, such as PacBio single-molecule real-time sequencing and Oxford Nanopore Technologies, generate reads spanning tens to hundreds of kilobases, enabling more accurate detection and precise characterization of large-scale genomic events (3), including complex rearrangements and repetitive regions (4, 5). These technological improvements have dramatically expanded the scope and resolution of SV detection, facilitating more comprehensive genomic analyses and enhancing our understanding of the functional and clinical impacts of genomic variation (6, 7). Nevertheless, a minority of SVs, particularly those spanning repetitive or poorly mappable regions, continue to exhibit imprecise or variable breakpoint placement across samples, with offsets that can reach hundreds of bases, highlighting that some technical challenges still persist (8).

Interpreting the functional consequences of SVs remains a major challenge as the mechanisms by which SVs exert phenotypic effects are highly heterogeneous and context-dependent. A single SV may simultaneously disrupt a gene, delete a regulatory element, and perturb chromatin architecture, making it difficult to predict the overall consequence using simple models. Traditional variant annotation tools, like CADD (9), were primarily developed with SNVs and small indels in mind, and they often fall short when applied to SVs. Therefore SV-specific tools have been developed to fill this gap, offering methods to prioritize, annotate, or filter SVs based on different biological features. Knowledge-based tools like AnnotSV (10) integrate gene, regulatory, and clinical annotations to highlight SVs overlapping known functional elements, making them useful for well-characterized regions and clinical variants. Other approaches, such as SVScore (11), estimate SV deleteriousness by aggregating CADD scores across the variant interval. While practical, these methods rely heavily on existing annotations or SNV-based models and may miss novel or context-dependent SV effects. More recent approaches use data-driven and machine learning methods to prioritize SVs based on learned relationships between genomic annotations and variant impact (12–14). Approaches like PhenoSV and POSTRE extend SV prioritization through phenotype-aware or cell-type-specific scoring (15, 16). Tools such as CADD-SV (12) and TADA (13) integrate diverse SV-level and genomic context features into unified scores, enabling prioritization beyond well-annotated regions or simple SNV-score aggregation. CADD-SV introduced a new training strategy by contrasting evolutionarily fixed SVs as proxy-neutral examples with simulated SVs as proxy-deleterious, avoiding reliance on clinical labels and less affected by clinical ascertainment bias. However, CADD-SV employed separate models for different SV types and contexts and was limited to coordinate-based annotations, constraining its generalizability and downstream flexibility.

Building upon these foundations, we present CADD-SV v2.0, a redesigned and enhanced framework for scoring the potential deleteriousness of structural variants. The new version introduces major changes and improvements by incorporating all supported SV types in a single model, expanding the training set substantially, and using new, updated and extended annotations for scoring SV effects. Further, with the advent of large language DNA models, we explore the incorporation of features directly derived from the altered sequence and its immediate context.

## Methods

### Source Data and Preprocessing

To construct the training dataset, we followed an evolutionary approach that contrasts naturally occurring structural variants (SVs) with simulated variants. Naturally occurring SVs that have persisted in human or non-human primate lineages were considered proxy-neutral, based on the assumption that variants tolerated over evolutionary timescales are less likely to have strongly deleterious effects. In contrast, proxy-deleterious SVs were generated separately by randomly placing variants in the human reference genome, using some matching constraints (See next paragraph). These simulated SVs have not been subject to the same evolutionary selection and are expected to include a greater proportion of functionally deleterious variants, although individual simulated variants are not necessarily pathogenic.

Proxy-neutral SVs were obtained from a previously published callset based on multispecies whole-genome alignments and structural variant detection (Kronenberg et al. 2018). The dataset included variants from humans, chimpanzees, gorillas, and orangutans. Fixed variants in non-human primates were defined as SVs with an allele frequency (AF) of at least 50% in at least one ape genome and absent from humans, whereas human-fixed variants had an AF of at least 50% in humans and were absent from all ape genomes. Only autosomal variants were retained. The resulting training dataset comprised 280,236 SVs, equally divided between proxy-neutral and proxy-deleterious classes. From this number we removed chromosome 8 to be used as a holdout set. A detailed summary of training and holdout set numbers are available in Supplementary Table 1. The training dataset and the code used for its generation are available in the Supplementary Files.

### Random Variant Generation

Proxy-deleterious variants consisted of randomly generated SVs in the human genome (GRCh38), matched by length and type to the proxy-neutral variants. Variants were randomly placed within chromosomes using bedtools shuffle, excluding regions of proxy-neutral variants, and constrained to ape-alignable genomic regions. Ape-alignable regions were identified by merging alignable segments between human and ape genomes from the original Kronenberg et al. (2018) callset, forming a continuous genomic background for randomization. To preserve the interpretability of these proxy-deleterious variants, additional filtering steps were applied to ensure they did not overlap annotated assembly gaps from UCSC (18), which are prone to spurious calls and low mapping quality (see Supplementary Files).

### Sequence Integration in Training Data

Sequences for insertion variants were integrated into the training dataset. Inserted sequences for proxy-neutral SVs were extracted from species-specific assemblies: Chimpanzee (GCA_002880755), Gorilla (GCA_900006655), Orangutan (GCA_002880775), Human cell line CHM13 (GCA_002884485), and Human sample YriHSA from the Yoruba population (GCA_002884475), using the respective target coordinates provided in the Kronenberg et al. 2018 VCF file. For proxy-deleterious insertions, random length-matched DNA segments from GRCh38 were sampled and included in the training feature set (see Supplementary Files).

### Feature Annotation and Extraction

Genomic features used for training were derived from publicly available annotation resources and encompassed conservation scores, overlaps with regulatory regions, proximity to genes, repeat content, and sequence composition metrics. Feature extraction involved a combination of bedtools v.2.27.1 (19), tabix from htslib v.1.10.2 (20), pysam v.0.21 (21), and custom scripts. For each variant, features were computed not only for the affected interval but also for 100 bp flanking windows around the affected interval, capturing the broader genomic environment. A comprehensive list of all features and their source is provided in Supplementary Table 2.

### Model Architecture and Training

A unified Random Forest classifier was trained to score all SV types, replacing four separate models used in CADD-SV v1.1. The classifier was implemented using the scikit-learn library (22) with the following parameters chosen using grid search evaluated on chr8 that was used as a holdout from the training set: n_estimators=500, max_depth=10, max_leaf_nodes=300, min_samples_split=2, min_samples_leaf=5, the rest was kept as default in the RandomForestClassifier function, despite a small increase in performance at n_estimators=1000 for the default model, we decided to retain the model with 500 estimators for size and performance reasons (Supplementary Fig. 1). To handle class imbalance, a class-weighting scheme was used during training to prioritize insertions, which were underrepresented relative to deletions. Sample weight was set to 1 for deletions and to the imbalance ratio (n_deletions/n_insertions) for insertions to reduce bias towards deletion-specific features, the training script is available in the github repository of CADD-SV.

### Sequence-Based Feature Extraction Using SegmentNT

To capture the functional genomic effects of structural variants (SVs) arising from changes in DNA sequence content and organization, we employed SegmentNT (23), a pretrained deep-learning model that predicts functional genomic elements directly from DNA sequences. SegmentNT generates nucleotide-resolution predictions for 14 functional classes, including genes, splice sites, promoters, enhancers, and other regulatory elements. Unlike static genomic annotations, these predictions depend on the local sequence context and can therefore reflect changes introduced by SVs, including those affecting multiple coding or non-coding elements. SegmentNT was selected for its efficiency on relatively short sequence windows and its functional element classification objective, which is well suited to the annotation-based framework of CADD-SV.

For each SV, we constructed reference (REF) and alternative (ALT) allele sequences representing the genomic context before and after the structural alteration, respectively. Sequence inputs were standardized to a maximum length of 1,200 bp, comprising a central region of up to 1,008 bp flanked by 96 bp of reference sequence on either side. These dimensions were selected to accommodate the non-overlapping 6-mer tokenization used by the underlying Nucleotide Transformer v2 (NTv2) model. The 96-bp flanking regions approximate the 100-bp flanks used for coordinate-based feature extraction while maintaining compatibility with the tokenization scheme. The 1,008-bp central region accommodates 79.34% of the SVs in the training dataset, providing a balance between sequence coverage and computational requirements. A detailed summary of training set size distribution is available in Supplementary Table 3.

Variant-specific REF and ALT sequences were generated using custom scripts. For insertions, the ALT sequence was constructed by incorporating the inserted sequence between the upstream and downstream reference flanks. For deletions, the deleted segment was removed and the remaining flanking sequences were joined. For duplications, an additional copy of the affected segment was inserted adjacent to its original location, whereas for inversions, the affected sequence was reverse-complemented while retaining the surrounding genomic context. For variants exceeding the 1,008-bp central-region limit, the central portion was truncated and represented using two placeholder N tokens, preserving the sequences adjacent to the variant boundaries. All sequences were generated in a strand-aware manner and stored in a standardized BED-like format.

The resulting REF and ALT sequences were processed separately using SegmentNT following the model documentation (https://huggingface.co/InstaDeepAI/segment_nt). For each sequence, SegmentNT generated nucleotide-level probabilities for all 14 functional classes. Predictions were stored in structured HDF5 files, and sequences were processed sequentially to limit memory consumption.

To derive interpretable sequence-based features, SegmentNT predictions were summarized separately for two regions: the complete input sequence, including the flanking regions, and the central region affected by the SV. For each region and functional class, three summary statistics were calculated: the mean, minimum, and maximum predicted probabilities. This procedure yielded 84 features per sequence (14 functional classes × 3 statistics × 2 regions), with REF and ALT features stored in separate tab-separated files.

We additionally quantified SV-induced changes in SegmentNT predictions within the flanking regions by comparing corresponding positions in REF and ALT sequences. For each functional class, predictions from the first and last 96 nucleotide positions were extracted, and absolute differences between REF and ALT probabilities were calculated position-wise. Because these flanking regions contain identical reference sequences in both representations, differences in their predicted probabilities capture the influence of the altered sequence context rather than changes in the underlying flanking nucleotide sequence. The resulting absolute differences were summarized using the mean, minimum, and maximum for each functional class, yielding 42 additional features per SV (14 functional classes × 3 statistics). These features provide a complementary measure of the predicted functional consequences of SVs on their surrounding genomic context.

### Validation Sets

To evaluate CADD-SV v2 performance we generated different validation sets:

-**ClinVar validation set:** SVs were derived from the ClinVar database (downloaded Sep 28, 2025 from the NCBI FTP server) (24). Variants mapped to the GRCh38 assembly and classified as non-single-nucleotide variants were retained. Pathogenic subset was extracted based on the “Clinical significance” field being either “Pathogenic” or “Likely pathogenic”, retaining only entries with established review criteria (“no assertion criteria provided” entries were excluded). Separate files were created for deletions (n=4346), duplications (n=364), insertions (n=403), and inversions (n=13) based on the “Type” field. SVs shorter than 50 bp were excluded to ensure consistency with the CADD-SV size definition. Coordinates for each variant type were formatted into BED files, with insertions including the sequence field when available. Each record was annotated with its corresponding SV type (DEL, DUP, INV, or INS) to produce the final ClinVar pathogenic BED files used for downstream validation.
-**gnomAD validation set:** Structural variants were obtained from the gnomAD-SV v4.1 dataset, mapped to the GRCh38 reference genome (1). To construct a benign validation set, common variants with intermediate allele frequencies (10–90%) were selected, representing polymorphic variants likely to be functionally neutral, separate files were created for deletions (n=7050), duplications (n=5164), insertions (n=5019), and inversions (n=19). For the deleterious set, rare variants were extracted by selecting events with extremely low population frequencies, excluding singletons to avoid sequencing artifacts, given the great amount of genomes that gnomAD-SV provides. Specifically, variants with an allele count of 2 out of more than 126,000 alleles were retained and separate files were created for deletions (n=44328), duplications (n=28769), insertions (n=6180), and inversions (n=397).
-**1KGP ONT validation set:** An additional validation set was derived from Oxford Nanopore long-read sequencing data generated by Schloissnig et al. (25), to include some sequence-resolved insertions in our validation. Structural variants were extracted from the SVs identified using the svim-asm pipeline and mapped to the GRCh38 reference assembly (Data downloaded on Sep 30, 2025 from: https://ftp.1000genomes.ebi.ac.uk/vol1/ftp/data_collections/1KG_ONT_VIENNA/release/ v1.1/svim-asm-hg38/). To approximate neutral and deleterious variant sets, allele frequency thresholds were applied to the population-level calls. Variants with approximately 10% minimum allele frequency were considered neutral, while rare variants occurring at or below 0.1% were classified as deleterious proxies. Separate files were generated for insertions (proxy-neutral n=21361, proxy-deleterious n=24648) and deletions (proxy-neutral n=12045, proxy-deleterious n=19396).
-**UK Biobank (deCODE) validation set:** Structural variant associations were obtained from the deCODE analysis of the UK Biobank cohort (Data downloaded on Sep 30, 2025 from: https://www.decode.com/ukbsummary/), based on ICD-10 phenotype summary statistics (26). The downloaded dataset contained genome-wide association results for structural variants tested across a wide range of phenotypes. To generate a high-confidence validation set, only variants with high imputation quality (INFO > 0.9), strong association signals (p-value < 0.001) and low allele frequency (AF < 0.001) were retained. Variants were parsed and converted into BED format by extracting chromosomal coordinates, inferring structural variant type (INS, DEL, DUP) from the effect allele annotation, and estimating variant length when size information was available. For end coordinate calculation, insertions were assigned a 1 bp span, while other SV types used reported or inferred size values. Phenotypes corresponding to congenital disorders were specifically selected to enrich for potentially pathogenic variants, while phenotypes in ICD-10 chapters A and B (infectious and parasitic diseases) were excluded. This filtering strategy produced a high-quality set of structural variants associated with congenital phenotypes, serving as an association based proxy for pathogenic SVs rather than a set of confirmed causal variants. Separate files were generated for deletions (n=946), duplications (n=31) and insertions (n=285).

Each deleterious set is then matched with neutral sets from gnomAD or 1000 Genomes, the larger set is then sampled to match the sample size of the smaller set. The validation sets used to generate the benchmarking plots are available in the Supplementary Files.

### Benchmarking

CADD-SV v2.0 was benchmarked against the following state-of-the-art models:

**CADD-SV v1.1:** Downloaded from its GitHub repository and run with default settings (12).

**AnnotSV v3.5.2:** Run through its web interface using default settings (27).

**TADA v1.0.2:** Installed through pip and run with default settings (13). Coordinates of validation sets were converted to hg19 (GRCh37) using liftOver with 60% minimum match (18). As TADA operates on a different reference genome assembly, this coordinate conversion may introduce minor differences in variant representation and consequently affect the benchmarking results.

Not included in the benchmark because already benchmarked against CADD-SV v1.1 in Kleinert & Kircher 2022: SVScore (11), TAD-fusion (28) and StrVCTVRE (29).

Two single-feature baselines were extracted directly from the CADD-SV annotation tables. The conservation baseline (PhastCons100_sum) was calculated by summing available 100-way vertebrate PhastCons scores across the SV interval (30). The gene-distance baseline (gene_dist) represents the minimum distance between the SV interval and an annotated gene interval from Ensembl release 96 (31), calculated using bedtools closest. Overlapping intervals were assigned a distance of zero. Both features underwent the standard CADD-SV transformation, log₁₀(|x| + 1), and were evaluated individually without classifier training or inclusion of flank features. Baseline AUROCs were reported independently of score direction as max(AUROC, 1 - AUROC).

### Model performance evaluation

Model performance was evaluated using the area under the receiver operating characteristic curve (AUROC). AUROC values were computed from continuous deleteriousness scores using roc_auc_score function from the scikit-learn library v.1.5.2 (22). Data handling and preprocessing were performed using pandas v.2.2.3 (32, 33) and NumPy v.2.1.3 (34), and model predictions were generated from pretrained classifiers serialized with joblib v.1.4.2 (35). Figures were produced using matplotlib v.3.10.8 (36). Uncertainty in AUROC estimates was assessed using 1,000 stratified bootstrap resamples for each validation dataset, SV type and model. Deleterious and neutral records were sampled separately with replacement, preserving the original number of records in each class. AUROC was recalculated for each resample, and 95% confidence intervals were defined by the 2.5th and 97.5th percentiles of the bootstrap distribution. Bar heights represent AUROCs calculated from the original validation records, rather than bootstrap means. All analyses were carried out in Python using these libraries.

### Snakemake Pipeline

All processing and modeling workflows were developed in Python and bash, and coordinated through a modular Snakemake (v7.31.0) pipeline (37). This pipeline integrates sequence extraction, SegmentNT inference, feature summarization, and variant annotation, enabling reproducible and scalable handling of large SV datasets. A dedicated training mode allows streamlined model generation using alternative training datasets. Logging, resource allocation, and job dependencies were defined at each stage. The code is available at https://github.com/kircherlab/CADD-SV.

### PHRED-Scaling

Raw variant scores were converted to PHRED-scaled values using a rank-based transformation. For each SV type, PHRED scaling was derived from a background set of 100,000 randomly generated variants. Scores were sorted in descending order such that larger scores corresponded to more extreme (deleterious) variants. Variants were assigned ranks r = 1, 2, …, N, and PHRED scores were computed as −10 × log10(r / N). To avoid unstable PHRED levels driven by single extreme observations, PHRED levels were only retained when supported by at least three variants; accordingly, ranks r < 3 were excluded, therefore the maximum PHRED score corresponds to 45. PHRED values were formatted using an extended PHRED scale: values ≥30 were rounded to the nearest integer, values between 20 and 30 were reported with one decimal place, values between 10 and 20 with two decimal places, and values below 10 with three decimal places, following conventions used in CADD’s PHRED-based scoring scheme (38). For each PHRED level, the cutoff score was defined as the least extreme score (i.e., the last-ranked value) mapping to that PHRED level. For each SV type, lookup tables for converting raw scores into PHRED-scaled values were generated. Importantly, the CADD-SV PHRED score represents the relative extremeness of an SV within a background distribution and should not be interpreted as a calibrated probability of pathogenicity.

## Results

### New and extended training set definition

CADD-SV v2.0 introduces a substantially improved training framework and model architecture. One of the most persistent challenges in the development of SV impact prediction models is the lack of sufficiently large, diverse, and unbiased training datasets. While resources such as ClinVar and the Human Gene Mutation Database (HGMD) offer curated annotations for SNVs and small indels, their coverage of structural variants remains extremely limited. The relatively small number of SVs available in ClinVar are typically pathogenic events, large in genomic size, coding-sequence altering, and located in or near well-studied genes. This strong bias toward clinically relevant and highly deleterious variants means they do not represent the broader spectrum of SVs observed in population-scale studies. As a result, they introduce significant ascertainment and selection biases, limiting their use as training data in models designed to generalize beyond known pathogenic genes and across molecular effects.

To address this issue, CADD-SV adopted an evolutionary approach to training data generation (Fig. 1A). In CADD-SV v1.1, proxy-neutral SVs were defined as variants that are fixed either on the human or chimpanzee lineage. In contrast, proxy-deleterious SVs were randomly sampled from the human genome, matched in type and length to their neutral counterparts. This contrast between proxy-neutral and randomly sampled, proxy-deleterious, variants allows the model to learn features associated with functional constraint without relying on incomplete or biased clinical annotations (See Methods).

In CADD-SV v2.0, this strategy is extended in several important ways to improve the size, diversity, and robustness of the training set. The updated proxy-neutral set now includes variants not only from humans and chimpanzees, but also from gorillas and orangutans (Fig. 1A). This broader phylogenetic sampling increases the evolutionary depth of the training data, capturing a more diverse set of tolerated SVs across the great ape lineage. Each species represents an independent lineage subjected to purifying selection, and their inclusion helps reduce potential lineage-specific artifacts. The proxy-deleterious set continues to be drawn from randomly generated SVs in the human genome (limited to alignable sequence between the ape species), matched in size and type to their neutral counterparts to maintain a meaningful and balanced contrast.

The total number of structural variants used for training in CADD-SV v2.0 is 265,117, comprising 50,379 insertions and 214,738 deletions. These are evenly divided between the proxy-neutral and proxy-deleterious classes. This marks a substantial increase in training data compared to CADD-SV v1.1, which relied on four separate models, each trained on a limited subset of variants from either humans or chimpanzees. In v1.1, the full training set across all four models included less than 80,000 SVs. Proxy-neutral sets consisted of 9,431 human deletions, 10,836 human insertions, 8,061 chimpanzee deletions, and 10,771 chimpanzee insertions (12). In contrast, the unified dataset used in v2.0 represents nearly four times the number of training examples, all used to train one single model instead of using different subsets of the total number to train different models. This increase in scale provides greater statistical power for the model to learn discriminative patterns of SV impact. Supplementary Fig. 2 shows the increase in performance on the holdout chromosome given by larger training sets. Deletion performance largely reached a plateau by approximately 100,000-150,000 training records, while insertion performance continued to show modest gains at larger sizes.

**Figure 1:**
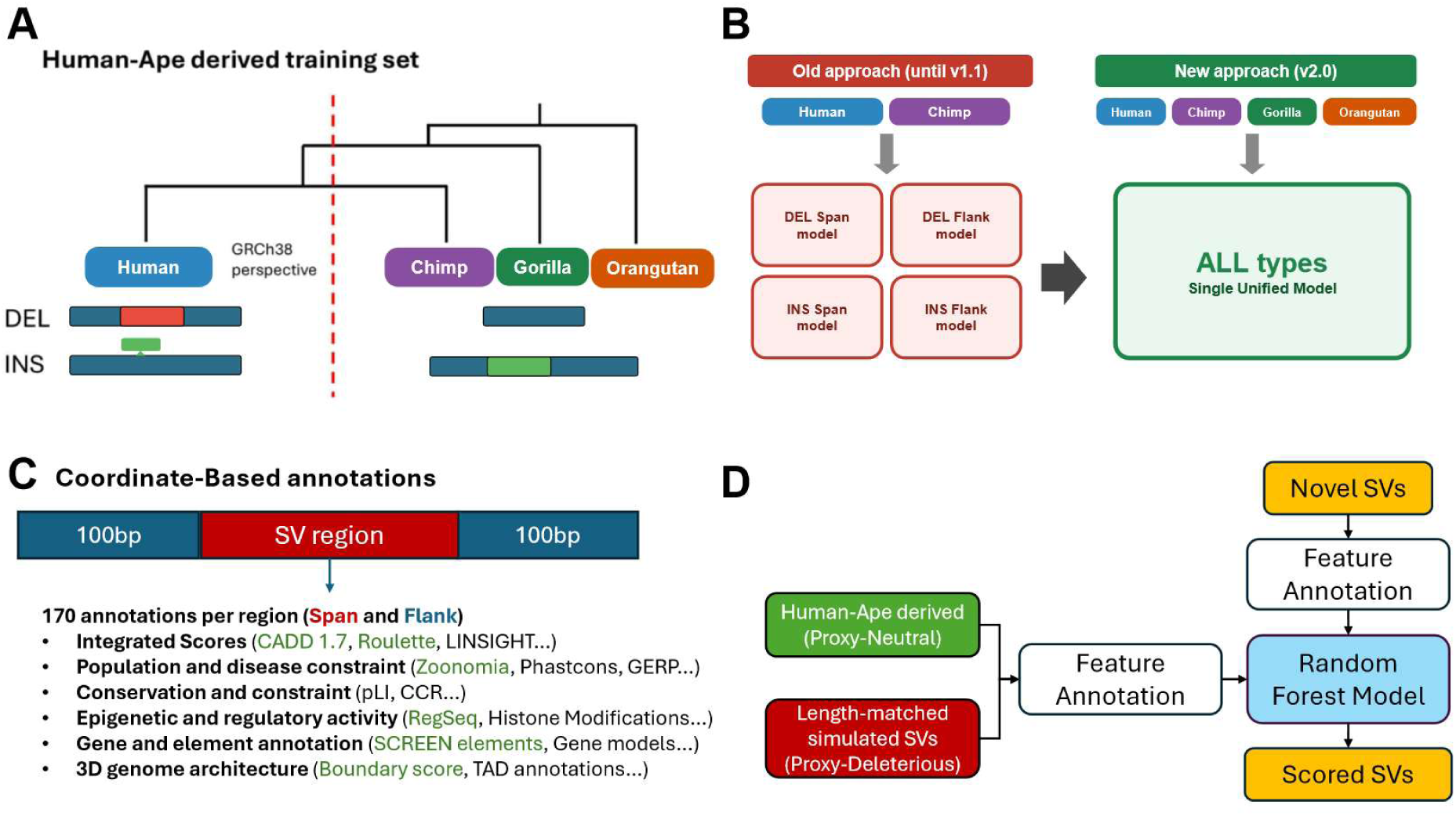
Overview figure. A) Human-ape derived training set construction. Proxy-neutral SVs are identified from variants fixed in great ape species or human and projected onto the human GRCh38 reference. B) Comparison of modeling strategies. The previous framework trained separate models for deletion span, deletion flank, insertion span, and insertion flank tasks using only human and chimpanzee data, whereas the updated approach integrates human, chimpanzee, gorilla, and orangutan variants into a single unified model that jointly supports all SV types. C) Coordinate-based feature annotation. Each SV is represented by the variant interval including 100bp flanking regions on both sides and annotated with 170 features spanning different categories. Highlighted in green the newly added annotations to v2.0. D) Diagram of CADD-SV model architecture.

### One Random Forest model for all SV types and an extended feature set

In the previous CADD-SV version, insertions and deletions were each modeled separately in two genomic contexts: the span of the SV and its surrounding flanks. The final impact score for a given variant used the maximum score from these two specialized insertion and deletion models, while duplications were approximated by using a different combination of two models (Deletion Span and Insertion Flank). In v2.0, combining insertions and deletions into a single model instead of keeping them separate (Fig. 1B), allows the model to focus on features that generalize across types. As a result, the model should be less sensitive to type-specific patterns and better suited to also score SV classes not included in training, such as duplications or inversions.

Increases in training set size allow us to add new coordinate-based features without a high risk of overfitting to a small dataset. As a result, annotations on flanking regions are now treated as separate features within the same model, rather than training a separate model specifically for flanking regions. This approach doubles the total number of features but maintains the use of a single unified model, eliminating the need to choose between different scores. To complement the expanded training data and new model architecture, CADD-SV v2.0 incorporates a comprehensive update of genomic annotations used for feature extraction. These include the most recent versions of widely used integrated scores such as CADD v1.7 and its predicted regulatory features from the RegSeq CNN (39), Roulette (40), updated constraint metrics calculated by the Zoonomia project (41), refined regulatory element annotations including SCREEN elements (42), and 3D genome architecture data such as TAD boundary scores (43) (Fig. 1C).

Each SV is annotated based on both its span and the flanking regions (100 bp upstream and downstream), resulting in 170 features per region (Fig. 1C). This expanded and modernized annotation set provides a more accurate and detailed functional context for each variant and enhances the ability to identify potentially deleterious SVs. Even though duplications and inversions are not explicitly represented in the training set, the v2.0 model (Fig. 1D) shows strong performance on these variant classes (Fig. 2C-D). The unified architecture enables the model to generalize to duplications and potentially to more complex rearrangements, while maintaining or improving performance on deletions and insertions. A comparison between CADD-SV v2.0 and v1.1 approaches is available in Supplementary Table 4.

**Figure 2:**
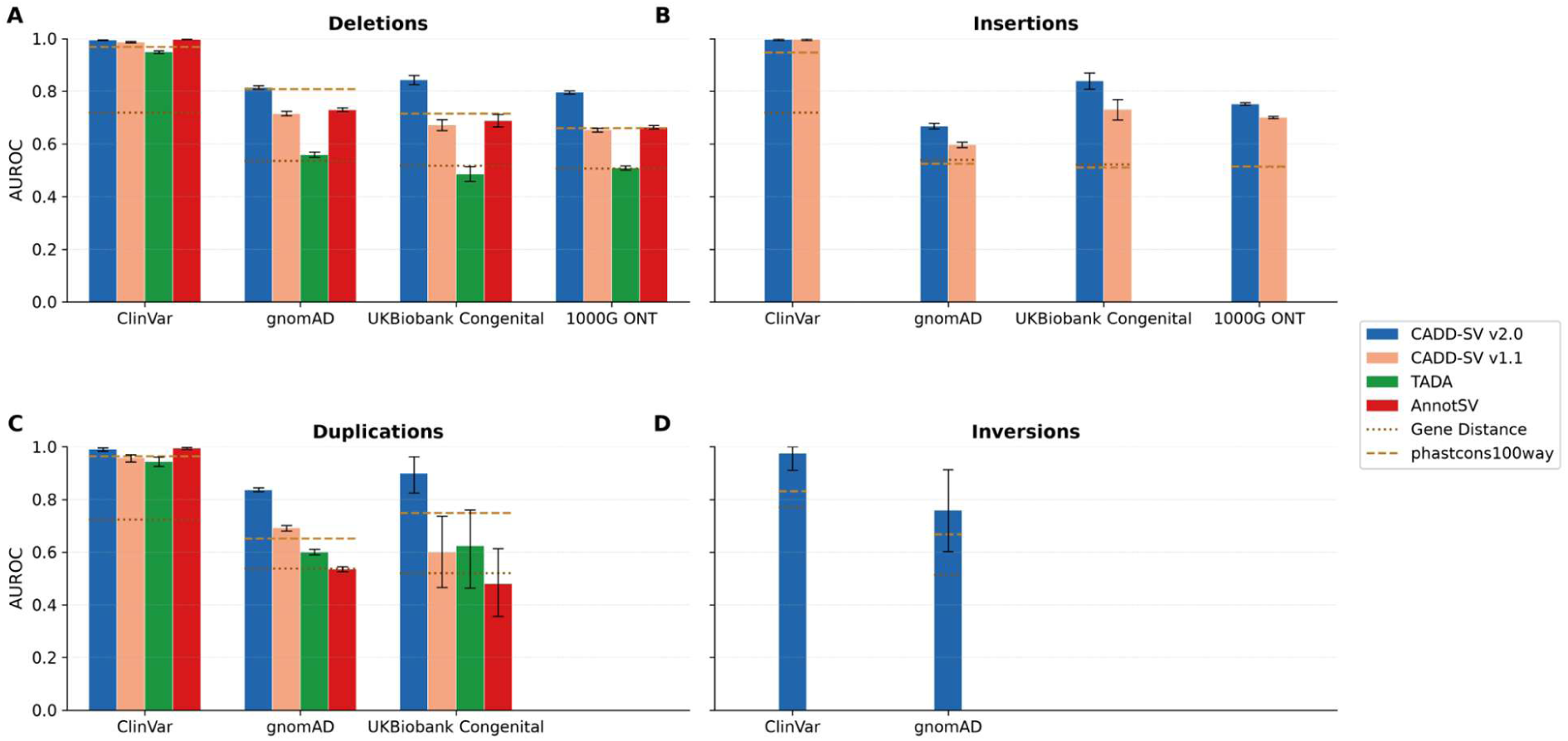
Comparative AUROC performance of structural variant deleteriousness scoring methods across independent validation datasets and variant classes. Bars show model performance for CADD-SV v2.0, CADD-SV v1.1, AnnotSV, and TADA; dotted and dashed horizontal lines indicate baseline predictors based on gene distance and PhastCons100way conservation. Error bars indicate 95% confidence intervals estimated from 1,000 stratified bootstrap resamples, as described in Methods. Panels summarize results for (A) deletions, (B) insertions, (C) duplications, and (D) inversions across ClinVar, gnomAD, UK Biobank congenital diseases, and 1000 Genomes ONT datasets (where available). Across nearly all benchmarks, CADD-SV v2.0 achieves the highest AUROC, with the largest gains observed in non-ClinVar datasets that are usually more challenging. Duplications have a substantial improvement over its previous version indicating that the new training and modeling strategy overcomes some of the biggest limitations of CADD-SV v1.1. Supplementary Fig. 3 shows AUROC as curves and values are shown in Supplementary Table 5, Precision-Recall and enrichment of Top scoring SVs are shown in Supplementary Fig. 4.

### Outperforming other SV scoring tools on an independent benchmark set

The combination of increased training data, broader evolutionary coverage, unified modeling, and richer annotations contributes to improved performance in SV impact prediction, the impact of each of these steps is shown in Supplementary Fig. 5, additionally Supplementary Figures 6-7 show impact of evolutionary features and training set weighting. Different validation sets were generated to evaluate the performance. The ClinVar validation set represents a high-confidence benchmark for pathogenic SVs, as it is based on expert-curated clinical annotations with established review criteria. This enables a reliable assessment of model sensitivity to disease-associated variants. However, the dataset is inherently biased toward well-characterized and clinically reported SVs, which may limit its representativeness of the broader spectrum of genomic variation. The gnomAD validation set provides a large-scale, population-based resource that supports robust evaluation across multiple SV types. The use of allele frequency thresholds allows for a systematic approximation of benign and deleterious variants, facilitating balanced benchmarking. Nevertheless, this classification strategy remains indirect, as rare variants are not necessarily deleterious or pathogenic and common variants are not always neutral, introducing a degree of label noise. The 1KGP ONT validation set contributes long-read, sequence-resolved SVs, improving representation of complex events and particularly enhancing the evaluation of insertions that are often underrepresented in short-read datasets. This allows benchmarking in a context that more closely reflects true genomic structure.

However, the reliance on allele frequency thresholds to define neutral and deleterious proxies reduces confidence in the assigned labels compared to clinically curated datasets. The UK Biobank (deCODE) validation set provides an independent cohort by selecting SVs statistically associated with congenital phenotypes, thereby offering a complementary evaluation framework rooted in genotype-phenotype relationships. In contrast to ClinVar, where pathogenicity is based on clinical curation, this approach identifies variants linked to disease in a population-scale, association-driven manner.

CADD-SV v2.0 outperforms its previous version, which was previously benchmarked with an older version of AnnotSV (10), SVScore (11), TAD-Fusion (28) and StrVCTVRE (29). It also shows stronger performance compared to other widely used tools in scoring structural variants such as TADA (13) and the latest version of AnnotSV (27), improving on all SV types (Fig. 2). In these benchmarks, we included gene distance and PhastCons100way baselines as the proximity to genes and evolutionary conservation are widely used indicators of deleteriousness, allowing us to assess whether CADD-SV v2.0 provides additional predictive value beyond these simple measures. Duplications (Fig. 2C) show a great increase in performance especially compared to CADD-SV v1.1, indicating that the new training and modeling approach was successful. As a practical benefit, the unified model supports scoring of SV types that can be represented within a single coordinate, including insertions, whereas some tools are limited in scope and unable to provide scores for certain variant classes. AnnotSV provides annotations for insertions but does not assign them a pathogenicity score. TADA, in contrast, is designed to prioritize copy-number variants (CNVs) and therefore supports deletions and duplications, but not insertions or inversions (Table 1). CADD-SV v2.0 extends scoring capabilities to inversions, a variant class not supported by most existing SV prioritization tools. Owing to the scarcity of well-annotated inversions in public resources, evaluation was limited to two validation sets derived from ClinVar and gnomAD. As shown in Figure 2D, CADD-SV demonstrates some performance improvement over baselines, however, these results should be interpreted with caution. The number of deleterious inversions is extremely limited (n = 13 in ClinVar and n = 19 in gnomAD), which substantially reduces statistical power. In addition, while the gnomAD-derived deleterious set follows the same frequency-based proxy approach used for other variant types, the very small number of inversions increases uncertainty and the potential impact of label noise. Furthermore, as inversions do not involve a net gain or loss of DNA sequence, they are, on average, more likely to be functionally neutral, which further complicates the identification of truly deleterious events. Consequently, these results should be considered preliminary and will require validation on larger and better-characterized inversion datasets.

**Table 1.**
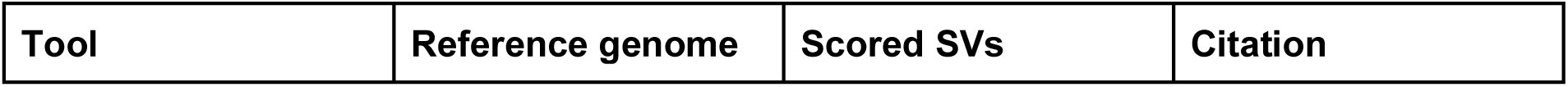
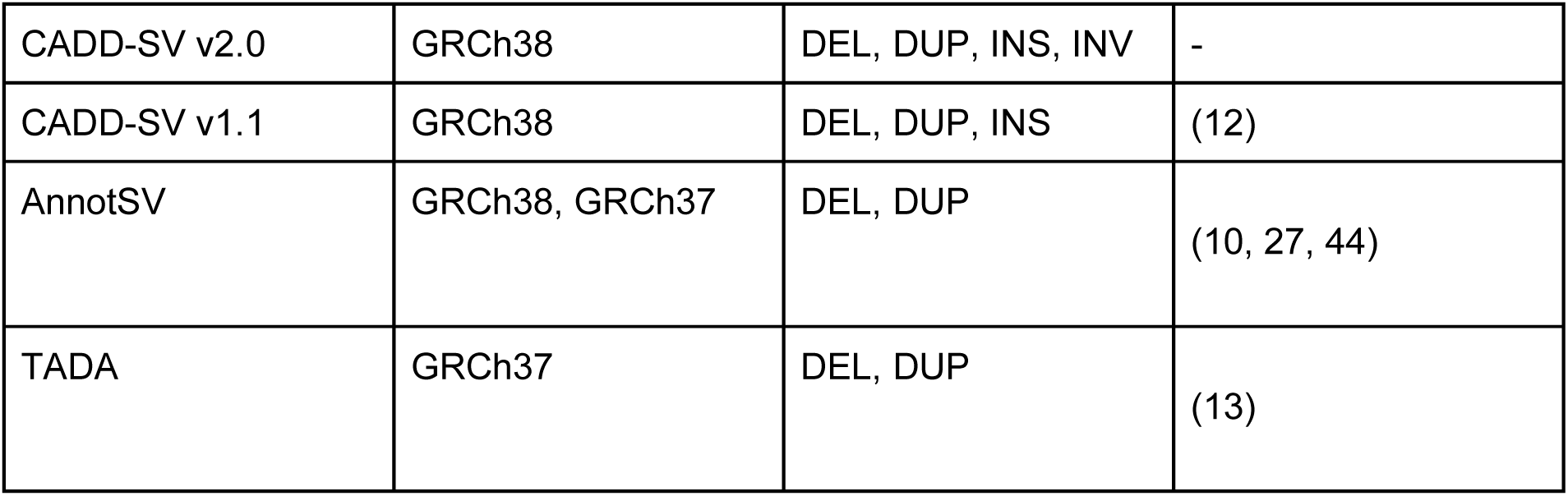
Table showing different reference genomes and scored SVs for the different tools used to benchmark CADD-SV v2.0.

### Sequence-Based Features and Integration of SegmentNT

To explore whether DNA sequence information could enhance feature representations and improve predictions of SV deleteriousness, we investigated the integration of sequence-derived annotations into our model. Given that sequence-resolved SVs represent an emerging and increasingly accessible class of genomic data, this approach potentially enables the model to capture information related to coding and non-coding element content, as well as sequence-level effects such as splice site disruption. However, incorporating such features requires access to sequence information for the training variants. For proxy-neutral insertions, we extracted inserted sequences from the original assemblies. For proxy-deleterious variants, we retrieved length-matched random DNA segments sampled from GRCh38, excluding specific regions as described in the Methods. This approach reflects that inserted sequences often derive from existing genomic material, making the reference genome a practical proxy for sequence composition, although it does not capture sequences absent from the reference (45) (See Methods).

**Figure 3:**
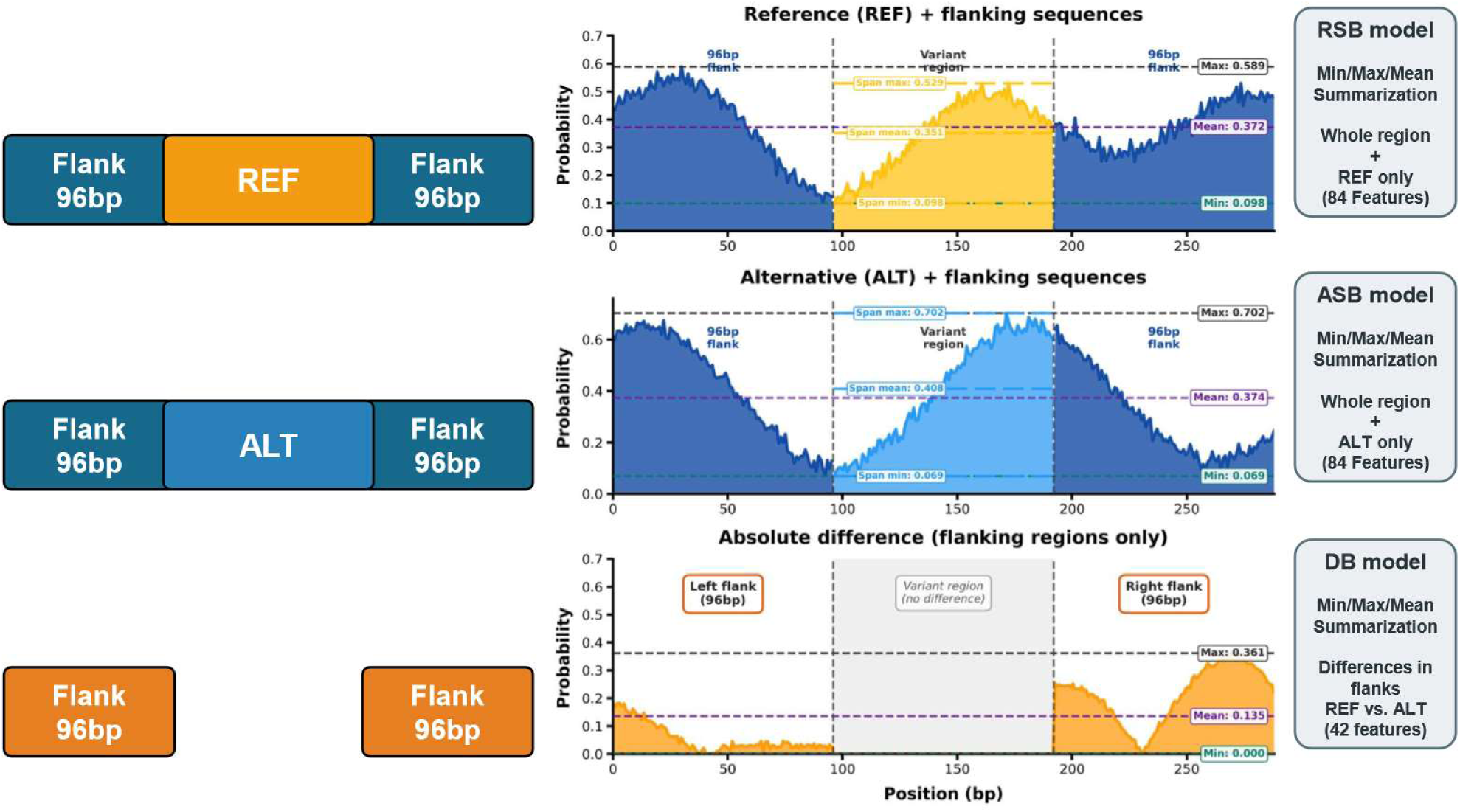
Overview of sequence-based feature extraction for variant effect modeling. A 96-bp flanking sequence is appended to both sides of the reference (REF) and alternative (ALT) alleles to generate two input windows of equal length. Predicted positional probabilities across the REF sequence (top) and ALT sequence (middle) are summarized using minimum, maximum, and mean values over the full region, and only the span region, to generate features for the reference sequence-based (RSB) and alternative sequence-based (ASB) models (84 features each). The absolute difference between REF and ALT predictions is then calculated for the flanking regions only, excluding the variant interval where the alleles differ (bottom), and summarized by minimum, maximum, and mean values to generate difference-based (DB) model features (42 features). Not shown in the figure: CADD-SV-SeqOnly model is the combination of all the features used for the RSB, ASB and DB models.

To transform the nucleotide-resolution SegmentNT output tracks into fixed-size numerical features, we computed summary statistics using multiple strategies. In the first approach, we derived reference sequence-based (RSB) and alternative sequence-based (ASB) features by calculating the mean, minimum, and maximum predicted probabilities for each functional class, separately across the entire input sequence and across the SV span alone (Fig. 3). These features were computed independently for the reference sequence (RSB) and the altered sequence (ASB), and corresponding models were trained using each feature set. In a second, more exploratory approach, we derived difference-based (DB) features by quantifying changes in functional predictions through differences in flanking-region probabilities between the reference and altered sequences. Because the flanking regions are identical in both inputs, this strategy isolates the effect of the sequence alteration on local functional predictions. These differences were likewise summarized using mean, minimum, and maximum statistics.

### Exploratory Evaluation of Sequence-Based Models

To assess the potential contribution of sequence-derived features to SV deleteriousness prediction, we performed an exploratory evaluation across multiple benchmark datasets. As shown in Figure 4, models relying exclusively on SegmentNT-derived features achieved consistent, albeit moderate, discrimination across ClinVar, gnomAD v4.1, UK Biobank Congenital, and 1000 Genomes ONT datasets. Sequence-only models showed discriminatory ability in several validation comparisons, with performance varying across SV types and datasets. Although they did not consistently outperform the coordinate-based baselines in every validation set, they enable scoring directly from sequences, extending applicability to non-reference haplotypes for which genomic annotation-based scores cannot readily be obtained. This result indicates that predictions based on raw DNA sequence alone capture signals relevant to SV deleteriousness.

**Figure 4:**
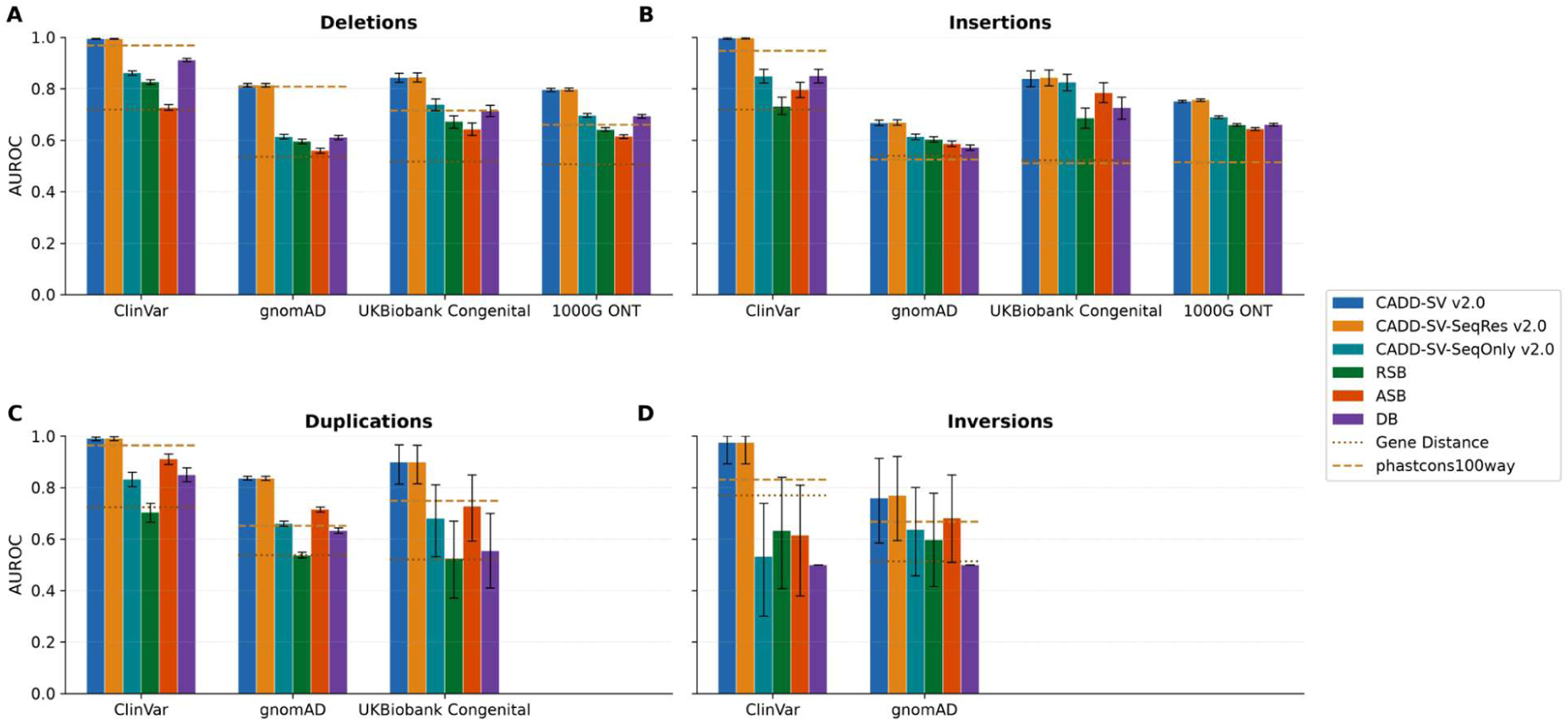
AUROC comparison of CADD-SV v2.0 and sequence-resolved derivative models across independent validation datasets and structural variant classes. Bars show performance for the full CADD-SV v2.0 model, CADD-SV-SeqRes v2.0, CADD-SV-SeqOnly v2.0, and component sequence-based models (RSB, ASB, DB); dotted/dashed horizontal lines indicate baseline predictors based on gene distance and PhastCons100way conservation. Error bars indicate 95% confidence intervals estimated from 1,000 stratified bootstrap resamples, as described in Methods. Panels summarize results for (A) deletions, (B) insertions, (C) duplications, and (D) inversions across ClinVar, gnomAD, UK Biobank congenital diseases, and 1000 Genomes ONT datasets (where available). Across nearly all benchmarks, the full CADD-SV v2.0 model and CADD-SV-SeqRes v2.0 consistently achieve the highest or tied-highest AUROC, demonstrating the value of coordinate-based and high-quality annotations. Integrating sequence-resolved features leads to modest improvement mainly on insertions. In contrast, the sequence-only models generally show lower and more variable performance than the full integrated models, but they frequently outperform simple baseline predictors, which could be particularly notable in settings where a richly annotated reference genome is unavailable. Supplementary Fig. 3 shows AUROC as curves and values are shown in Supplementary Table 5, Precision-Recall and enrichment of Top scoring SVs are shown in Supplementary Fig. 8.

Across datasets, the relative performance of DB, RSB and ASB varied by SV type. DB outperformed both RSB and ASB for deletions across all four validation datasets, whereas ASB performed better than DB for duplications and inversions. Notably, DB achieved competitive performance using only 42 features derived from reference-alternative differences across a limited 192-bp breakpoint context, compared with 84 features each for RSB and ASB. These results highlight the predictive value of local changes in functional potential and suggest that difference-based features can complement absolute sequence-content predictions. However, they do not by themselves establish that DB provides non-redundant information; this would require comparison of models trained with and without DB features. Breakpoint-context differences may also miss functional elements fully contained within an SV, such as an entire deleted gene or exon, supporting the rationale for combining complementary sequence representations in CADD-SV-SeqOnly. Performance at different SV sizes and genomic context (coding and non-coding) is shown in Supplementary Fig. 9.

When sequence-based features used for the DB model were integrated with the standard coordinate-based annotations in the CADD-SV-SeqRes model, only marginal improvements in overall performance were observed across most datasets (Fig. 4). In some benchmarks, performance gains were subtle and dataset-specific, while in others the combined model largely overlapped with the coordinate-based baseline. This limited improvement suggests that, although SegmentNT-derived features encode biologically meaningful information, they may be largely redundant or less informative than existing annotation-based features. Feature importance for the 3 models is shown in Supplementary Fig. 10, feature correlations and grouped permutation importance are shown for each model separately in Supplementary Figures 11-13.

## Discussion

The advancements introduced in CADD-SV v2.0 address several limitations in SV impact prediction, most notably the inadequacy of existing training datasets and the challenges of modeling a wide range of variant types. By expanding the proxy-neutral dataset to include fixed variants from multiple great ape lineages, we have substantially increased the size and diversity of the training data from less than 80,000 to more than 260,000 SVs. This change may reduce the influence of lineage-specific artifacts (i.e., due to genome assembly quality or the over-/underrepresentation of certain repeat elements (46)) and provide a more robust foundation for scoring variants. While the CADD approach originally introduced for short sequence variants (9) always dealt with several millions of variants in its Chimpanzee-derived training set, training set size was a concern for CADD-SV where only tens of thousand variants are identified from whole genome alignments. This leads to a much smaller training set and incomprehensive coverage of values across hundreds of features. It creates the risk of overfitting, especially when further annotations are added, but the new ape-derived training set considerably improves on that.

The decision to unify the training of insertions and deletions within a single model marks a key departure from the segmented approach of the previous release. While version 1 relied on four separate Random Forests and thus fragmented and small training sets, v2.0 consolidated training into a single model. Treating flanking annotations as part of a single model rather than building separate models for different regions enables more interpretable scoring while leveraging the full power of the dataset. These structural changes not only simplify the architecture but also enhance the model’s ability to generalize across SV types, including duplications, previously handled by different combinations of models, and an extension to inversions, although the performance should be interpreted with caution given the small size of the validation set. The unified model avoids the need to combine separate scores and is more consistent in its treatment of different variant classes.

Despite these improvements, some limitations remain. The main limitation of our training and benchmarking strategy is that proxy-deleterious variants are not necessarily truly deleterious, introducing noise into both training and validation. Although, this approach allows us to use large and diverse datasets while reducing the ascertainment biases associated with currently available clinically annotated variant datasets. CADD-SV cannot score SV types that cannot be represented as a single coordinate (i.e. Translocations). The integration of sequence-based features using SegmentNT demonstrated only marginal improvements in predictive performance, particularly for insertions and duplications. While the SegmentNT model provides base-pair resolution functional annotations from raw DNA, the additional computational cost of requiring a GPU to run the sequence models raises questions about its utility in standard workflows. We want to highlight though that the primary advantage of sequence-based approaches may emerge in settings where high-quality coordinate-based annotations are unavailable or incomplete. Here, sequence-only approaches could be useful. Such scenarios include analyses based on non-canonical references, telomere-to-telomere (T2T) assemblies (47), pangenome graphs (48), or personalized and population-specific genomes, where comprehensive functional annotations may lag behind sequence generation. In these contexts, sequence-based models offer a reference-agnostic mechanism to infer functional potential directly from DNA sequence, enabling SV impact assessment even in poorly annotated or novel genomic regions.

For these reasons, we are releasing the “CADD-SV-SeqRes” and “CADD-SV-SeqOnly” models as part of our computational pipeline, which otherwise uses the CADD-SV v2.0 model with coordinate-based annotations approach. The “CADD-SV-SeqRes” model uses the sequence derived features from the DB model in addition to coordinate-based annotations, which might give an advantage for sequence-resolved insertions. Further, “CADD-SV-SeqOnly”, a model that includes all the features generated from SegmentNT, is available as a reference-agnostic CADD-SV model, to be used in cases in which “REF” and “ALT” sequences are resolved and available but not anchored on the GRCh38 reference, allowing users to score SVs from personalized genomes, other species or germline vs. somatic alterations.

Overall, CADD-SV v2.0 demonstrates improvements in SV impact prediction by addressing key limitations in data, architecture, and annotation. The model’s increased predictive performance to distinguish between neutral and deleterious variants, along with broader SV type support, highlights its potential utility in both research and clinical settings regarding prioritization of SVs. At the same time, continued refinement of sequence modeling, better handling of complex SVs, and optimization of computational tradeoffs remain important areas for future development.

## Data Availability

The CADD-SV v2.0 source code, documentation, and workflow implementation are available from the CADD-SV GitHub repository at https://github.com/kircherlab/CADD-SV. CADD-SV can also be used through the online scoring webserver at https://kircherlab.github.io/CADD-SV/. The command-line package “caddsv” is available through Bioconda and PyPI. Supplementary Files associated with this study, including source code, notebooks, processed training and validation datasets are available from Zenodo at https://doi.org/10.5281/zenodo.23920326.

## Author Contributions

O.C. and M.K. conceived the study. O.C. developed the pipeline, generated training and validation datasets, performed model training, benchmarking, and analysis, and drafted the manuscript. M.K. supervised the study, contributed to study design and interpretation, and revised the manuscript. All authors approved the final manuscript.

## Acknowledgements

We thank Angelina Göbel-Knapp for her help with the new implementations on the CADD-SV webserver. We also thank current and previous members of the Kircher lab for helpful discussions and suggestions. Computational analyses were performed on the OMICS cluster at the University of Lübeck as well as on the HPC for Research cluster of the Berlin Institute of Health at Charité – Universitätsmedizin Berlin.

## References

1. Collins, R.L., Brand, H., Karczewski, K.J., et al. (2020) A structural variation reference for medical and population genetics. Nature, 581, 444–451.

2. Collins, R.L. and Talkowski, M.E. (2025) Diversity and consequences of structural variation in the human genome. Nat. Rev. Genet., 26, 443–462.

3. Mahmoud, M., Gobet, N., Cruz-Dávalos, D.I., et al. (2019) Structural variant calling: the long and the short of it. Genome Biol., 20, 246.

4. Sedlazeck, F.J., Rescheneder, P., Smolka, M.,> et al. (2018) Accurate detection of complex structural variations using single-molecule sequencing. Nat. Methods, 15, 461– 468.

5. Logsdon, G.A., Ebert, P., Audano, P.A., et al. (2025) Complex genetic variation in nearly complete human genomes. Nature, 10.1038/s41586-025-09140-6.

6. Chaisson, M.J.P., Sanders, A.D., Zhao, X., et al. (2019) Multi-platform discovery of haplotype-resolved structural variation in human genomes. Nat. Commun., 10, 1784.

7. Ahsan, M.U., Liu, Q., Perdomo, J.E., et al. (2023) A survey of algorithms for the detection of genomic structural variants from long-read sequencing data. Nat. Methods, 20, 1143–1158.

8. Audano, P.A. and Beck, C.R. (2024) Small polymorphisms are a source of ancestral bias in structural variant breakpoint placement. Genome Res., 34, 7–19.

9. Kircher, M., Witten, D.M., Jain, P., et al. (2014) A general framework for estimating the relative pathogenicity of human genetic variants. Nat. Genet., 46, 310–315.

10. Geoffroy, V., Herenger, Y., Kress, A., et al. (2018) AnnotSV: an integrated tool for structural variations annotation. Bioinforma. Oxf. Engl., 34, 3572–3574.

11. Ganel, L., Abel, H.J. and Hall, I.M. (2017) SVScore: an impact prediction tool for structural variation. Bioinformatics, 33, 1083–1085.

12. Kleinert, P. and Kircher, M. (2022) A framework to score the effects of structural variants in health and disease. Genome Res., 32, 766–777.

13. Hertzberg, J., Mundlos, S., Vingron, M., et al. (2022) TADA—a machine learning tool for functional annotation-based prioritisation of pathogenic CNVs. Genome Biol., 23, 67.

14. Lv, K., Chen, D., Xiong, D., et al. (2023) dbCNV: deleteriousness-based model to predict pathogenicity of copy number variations. BMC Genomics, 24, 131.

15. Sánchez-Gaya, V. and Rada-Iglesias, A. (2023) POSTRE: a tool to predict the pathological effects of human structural variants. Nucleic Acids Res., 51, e54.

16. Xu, Z., Li, Q., Marchionni, L., et al. (2023) PhenoSV: interpretable phenotype-aware model for the prioritization of genes affected by structural variants. Nat. Commun., 14, 7805.

17. Kronenberg, Z.N., Fiddes, I.T., Gordon, D., et al. (2018) High-resolution comparative analysis of great ape genomes. Science, 360.

18. Hinrichs, A.S., Karolchik, D., Baertsch, R., et al. (2006) The UCSC Genome Browser Database: update 2006. Nucleic Acids Res., 34, D590–D598.

19. Quinlan, A.R. and Hall, I.M. (2010) BEDTools: a flexible suite of utilities for comparing genomic features. Bioinformatics, 26, 841–842.

20. Li, H., Handsaker, B., Wysoker, A., et al. (2009) The Sequence Alignment/Map format and SAMtools. Bioinformatics, 25, 2078–2079.

21. Danecek, P., Bonfield, J.K., Liddle, J., et al. (2021) Twelve years of SAMtools and BCFtools. GigaScience, 10, giab008.

22. Pedregosa, F., Varoquaux, G., Gramfort, A., et al. (2018) Scikit-learn: Machine Learning in Python. 10.48550/arXiv.1201.0490.

23. de Almeida, B.P., Dalla-Torre, H., Richard, G., et al. (2025) Annotating the genome at single-nucleotide resolution with DNA foundation models. Nat. Methods, 22, 2301–2315.

24. Landrum, M.J., Lee, J.M., Riley, G.R., et al. (2014) ClinVar: public archive of relationships among sequence variation and human phenotype. Nucleic Acids Res., 42, D980–D985.

25. Schloissnig, S., Pani, S., Ebler, J., et al. (2025) Structural variation in 1,019 diverse humans based on long-read sequencing. Nature, 644, 442–452.

26. Halldorsson, B.V., Eggertsson, H.P., Moore, K.H.S., et al. (2022) The sequences of 150,119 genomes in the UK Biobank. Nature, 607, 732–740.

27. Geoffroy, V., Lamouche, J.-B., Guignard, T., et al. (2023) The AnnotSV webserver in 2023: updated visualization and ranking. Nucleic Acids Res., 51, W39–W45.

28. Huynh, L. and Hormozdiari, F. (2019) TAD fusion score: discovery and ranking the contribution of deletions to genome structure. Genome Biol., 20, 60.

29. Sharo, A.G., Hu, Z., Sunyaev, S.R., et al. (2022) StrVCTVRE: A supervised learning method to predict the pathogenicity of human genome structural variants. Am. J. Hum. Genet., 109, 195–209.

30. Siepel, A., Bejerano, G., Pedersen, J.S., et al. (2005) Evolutionarily conserved elements in vertebrate, insect, worm, and yeast genomes. Genome Res., 15, 1034– 1050.

31. Cunningham, F., Achuthan, P., Akanni, W., et al. (2019) Ensembl 2019. Nucleic Acids Res., 47, D745–D751.

32. The pandas development team (2020) pandas-dev/pandas: Pandas. 10.5281/zenodo.3509134.

33. McKinney, W. (2010) Data Structures for Statistical Computing in Python. In Walt, S. van der, Millman, J. (eds), Proceedings of the 9th Python in Science Conference.pp. 56– 61.

34. Harris, C.R., Millman, K.J., van der Walt, S.J., et al. (2020) Array programming with NumPy. Nature, 585, 357–362.

35. The joblib developers (2025) joblib. 10.5281/zenodo.14915601.

36. Hunter, J.D. (2007) Matplotlib: A 2D graphics environment. Comput. Sci. Eng., 9, 90–95.

37. Mölder, F., Jablonski, K.P., Letcher, B., et al. (2021) Sustainable data analysis with Snakemake. 10.12688/f1000research.29032.1.

38. Rentzsch, P., Witten, D., Cooper, G.M., et al. (2019) CADD: predicting the deleteriousness of variants throughout the human genome. Nucleic Acids Res., 47, D886–D894.

39. Schubach, M., Maass, T., Nazaretyan, L., et al. (2024) CADD v1.7: using protein language models, regulatory CNNs and other nucleotide-level scores to improve genome-wide variant predictions. Nucleic Acids Res., 52, D1143–D1154.

40. Seplyarskiy, V., Koch, E.M., Lee, D.J., et al. (2023) A mutation rate model at the basepair resolution identifies the mutagenic effect of polymerase III transcription. Nat. Genet., 55, 2235–2242.

41. Genereux, D.P., Serres, A., Armstrong, J., et al. (2020) A comparative genomics multitool for scientific discovery and conservation. Nature, 587, 240–245.

42. Moore, J.E., Purcaro, M.J., Pratt, H.E., et al. (2020) Expanded encyclopaedias of DNA elements in the human and mouse genomes. Nature, 583, 699–710.

43. Li, C., Bonder, M.J., Syed, S., et al. (2024) An integrative TAD catalog in lymphoblastoid cell lines discloses the functional impact of deletions and insertions in human genomes. Genome Res., 34, 2304–2318.

44. Geoffroy, V., Guignard, T., Kress, A., et al. (2021) AnnotSV and knotAnnotSV: a web server for human structural variations annotations, ranking and analysis. Nucleic Acids Res., 49, W21–W28.

45. Kehr, B., Helgadottir, A., Melsted, P., et al. (2017) Diversity in non-repetitive human sequences not found in the reference genome. Nat. Genet., 49, 588–593.

46. Yoo, D., Rhie, A., Hebbar, P., et al. (2025) Complete sequencing of ape genomes. Nature, 641, 401–418.

47. Nurk, S., Koren, S., Rhie, A., et al. (2022) The complete sequence of a human genome. Science, 376, 44–53.

48. Liao, W.-W., Asri, M., Ebler, J., et al. (2023) A draft human pangenome reference. Nature, 617, 312–324.

