## Supplementary Figures for "CADD-SV v2.0: An Extended Framework for Structural Variant Deleteriousness Prediction Using Coordinate- and Sequence-Based Features"

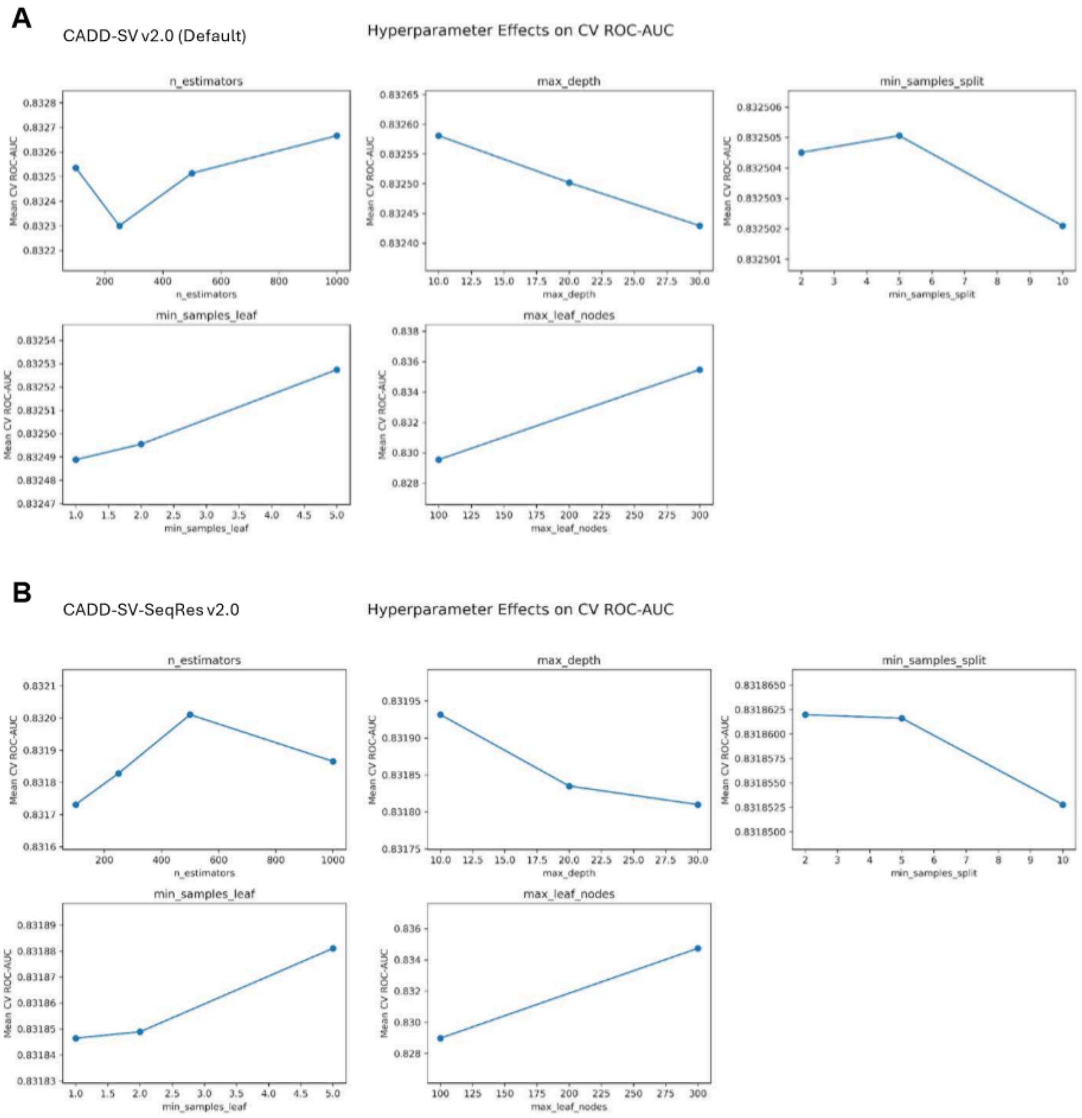


**Supplementary Figure 1: Sensitivity of model performance to hyperparameter settings for CADD-SV v2.0 variants.** Plots show the effect of key Random Forest hyperparameters on mean cross-validated ROC-AUC (y-axis) for two model configurations: ***(A)*** CADD-SV v2.0 (default feature set) and ***(B)*** CADD-SV-SeqRes v2.0. Each panel varies a single hyperparameter while holding others constant, including the number of trees (n_estimators), maximum tree depth (max_depth), maximum number of leaf nodes (max_leaf_nodes), and minimum samples required for node splitting (min_samples_split). Across both models, increasing the number of trees improves performance up to a plateau, consistent with ensemble stabilization. Tree complexity parameters show clear trade-offs: intermediate values of max_depth and max_leaf_nodes yield optimal ROC-AUC, whereas overly shallow trees underfit and overly complex trees show diminishing returns. Increasing min_samples_split reduces performance, indicating that allowing finer splits improves model flexibility and predictive accuracy. Based on these trends, final model hyperparameters were selected via grid search implemented in scikit-learn, using chromosome 8 as a holdout set separate from training data. The chosen configuration (n_estimators = 500, max_depth = 10, max_leaf_nodes = 300, min_samples_split = 2, min_samples_leaf = 5; all other parameters at default settings in RandomForestClassifier) reflects a balance between predictive performance and generalization. Despite a small improvement at 1000 n_estimators for the default model, we decided to retain the model with 500 n_estimator for size and performance reasons.

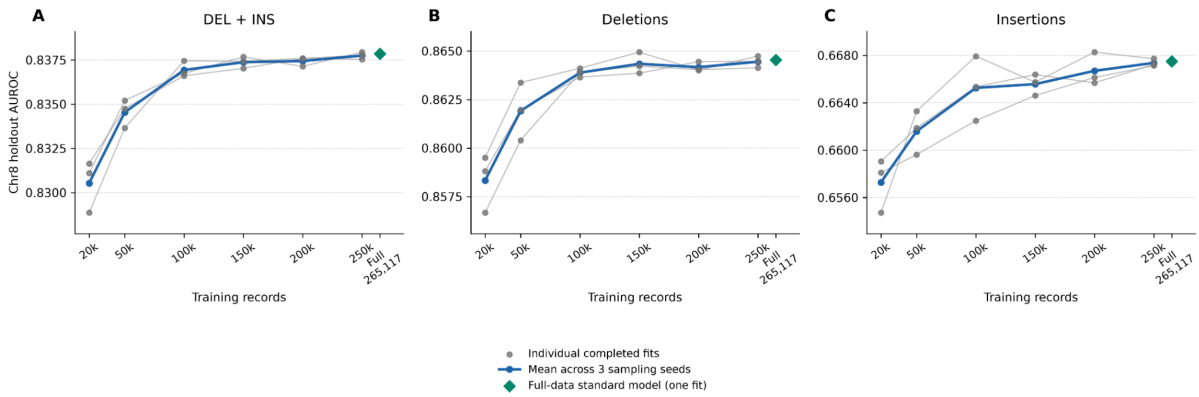


**Supplementary Figure 2. Effect of training-set size on CADD-SV v2.0 performance.** AUROC on the fixed chromosome 8 holdout is shown for ***(A)*** deletions and insertions together, ***(B)*** deletions and ***(C)*** insertions. The standard coordinate-based model was trained on subsets of 20,000, 50,000, 100,000, 150,000, 200,000 and 250,000 records. The 20,000-record subset approximates the training-set size of individual CADD-SV v1.1 models and serves as a size reference, not as a reconstruction of v1.1. Each subset contained equal numbers of proxy-deleterious and proxy-neutral variants. Subsets were nested within each of three sampling seeds. Features, random-forest hyperparameters and the sample-weighting rule were held constant across fits. Thin gray lines connect fits from the same sampling seed; the thick blue line shows the mean AUROC across seeds. The green diamond denotes the standard model trained on all 265,117 non-chromosome-8 records. AUROC generally increased with training-set size, with most of the pooled gain observed by 100,000 records and smaller gains thereafter. Panels use different y-axis scales.

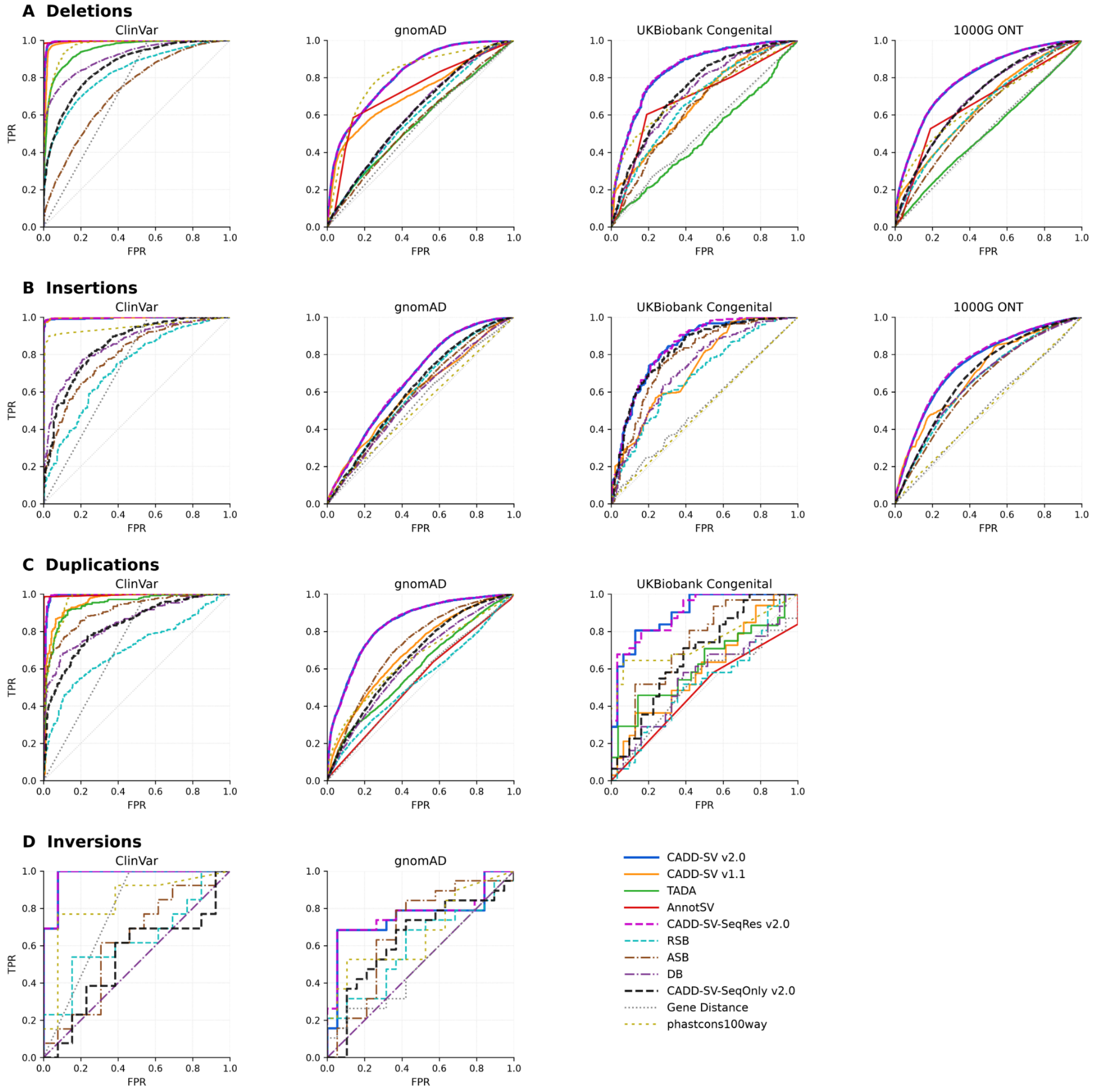


**Supplementary Figure 3: ROC-based comparison of structural variant deleteriousness scoring methods across validation datasets and variant classes.** Receiver operating characteristic (ROC) curves are shown for CADD-SV v2.0, CADD-SV-SeqRes v2.0, CADD-SV-SeqOnly v2.0, CADD-SV v1.1, AnnotSV, TADA, the sequence-based component models RSB, ASB and DB, and the gene-distance and PhastCons100way baselines. Rows correspond to ***(A)*** deletions, ***(B)*** insertions, ***(C)*** duplications and ***(D)*** inversions; columns correspond to ClinVar, gnomAD, UK Biobank congenital disease-associated variants and 1000 Genomes ONT variants. Comparisons are shown where data and method predictions are available. Curves depict true positive rate (TPR) against false positive rate (FPR); the diagonal indicates random discrimination. The gene-distance baseline is represented by a dotted gray line. Corresponding AUROC values, sample counts and 95% percentile confidence intervals calculated from 1,000 stratified bootstrap resamples are reported in Supplementary Table 5; confidence intervals were not calculated for the baseline predictors. CADD-SV v2.0 and CADD-SV-SeqRes v2.0 show comparable discrimination across most benchmarks, whereas sequence-only and component-model performance varies by SV class and dataset. Inversion comparisons are limited to ClinVar and gnomAD, with substantial uncertainty associated with the small evaluation sets.

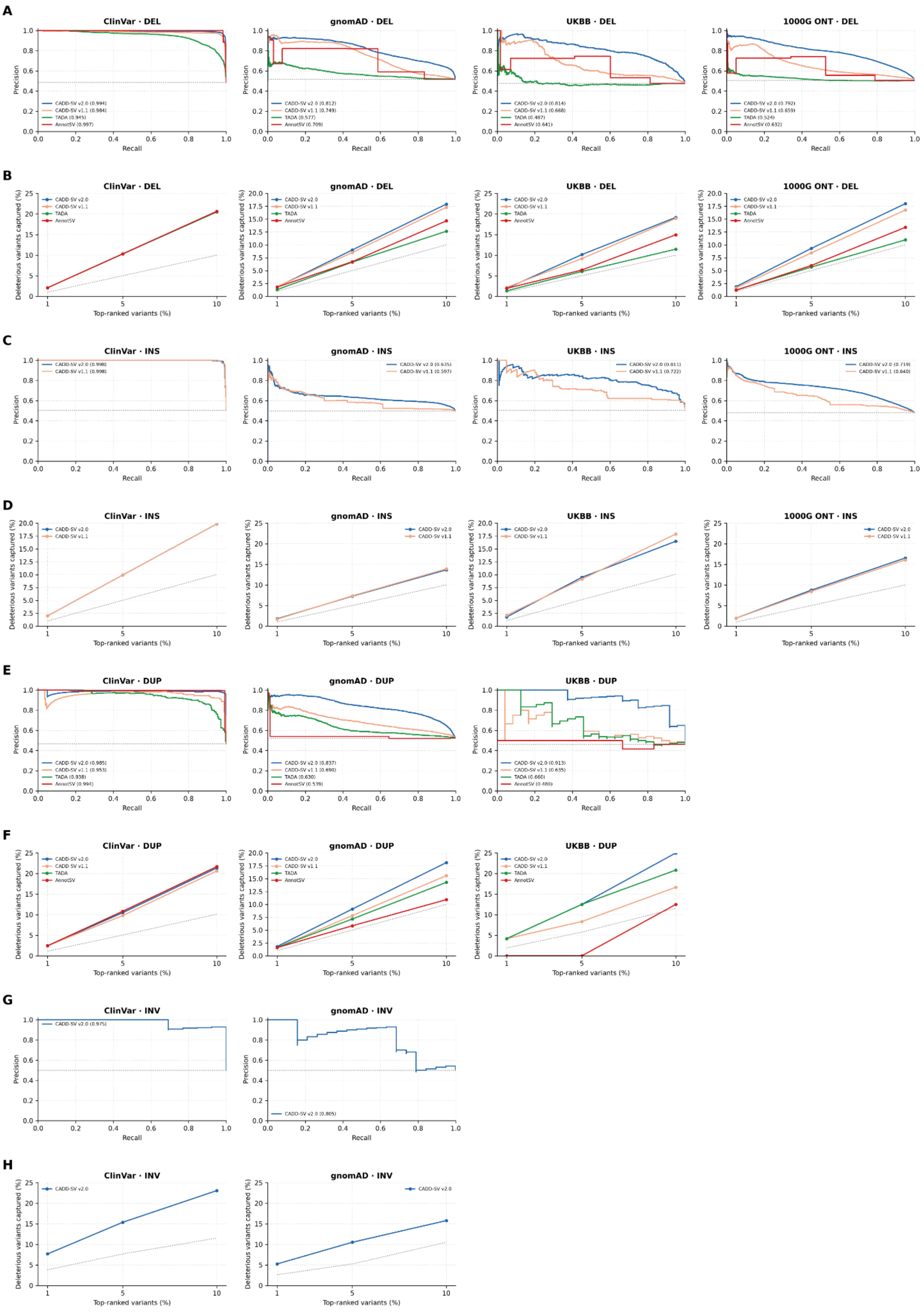


**Supplementary Figure 4: Precision-recall performance and enrichment of top-ranked structural variants for CADD-SV v2.0 and other methods.** Precision-recall curves (***A, C, E, G***) and the percentage of deleterious variants captured among the top-ranked 1%, 5%

and 10% of variants (***B, D, F, H***) are shown for deletions (***A, B***), insertions (***C, D***), duplications (***E, F***) and inversions (***G, H***). Columns correspond to ClinVar, gnomAD, UK Biobank congenital-disorder (UKBB) and 1000 Genomes ONT (1000G ONT) validation datasets, where available. Comparisons include CADD-SV v2.0, CADD-SV v1.1, TADA and AnnotSV, where results are available. Values reported in the precision-recall legends indicate average precision; dotted horizontal or diagonal reference lines indicate the dataset-specific baseline. CADD-SV v2.0 generally maintains strong precision across recall levels and captures a relatively high fraction of deleterious variants within the top-ranked groups, particularly for deletion and duplication benchmarks. Inversion results are restricted to the available ClinVar and gnomAD datasets.

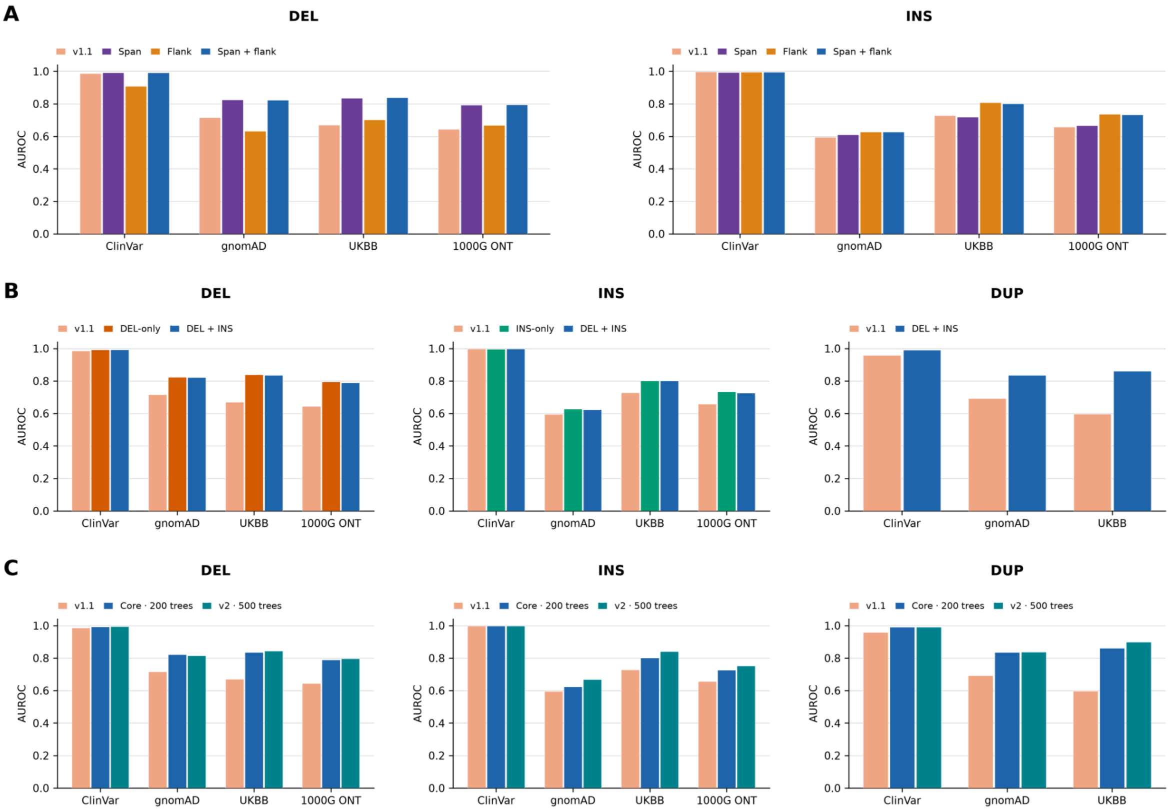


**Supplementary Figure 5: Contributions of genomic context, combined SV-type training and updated model configuration to CADD-SV performance.** Bars show the AUROC on the ClinVar, gnomAD, UK Biobank congenital-disorder (UKBB) and 1000 Genomes ONT (1000G ONT) validation sets, where available. CADD-SV v1.1 is included throughout as the reference. ***(A)*** Genomic context. Models trained separately on Deletions or Insertions are compared using span features, flank features or their combination. These models use the core annotation set and 200 trees, with 100-bp flanks. Within each SV-type comparison, the training subset and hyperparameters are held constant, allowing the contribution of span and flank information to be assessed. Span+Flank approach was chosen because it gives a single score, removing the need to choose between 2 different score as it was in v1.1. ***(B)*** Training SV-type composition. DEL-only and INS-only models are compared with a model trained jointly on DELs and INSs. All models in this comparison use core Span+Flank features. DEL validation includes DEL-only and pooled training; INS validation includes INS-only and pooled training. DUP validation evaluates transfer of the pooled model to an SV type not included in its training. ***(C)*** Updated v2 model configuration. The pooled core model, comprising v1.1 features, is compared with the v2 standard model, comprising the newly added features and revised random forest hyperparameters.

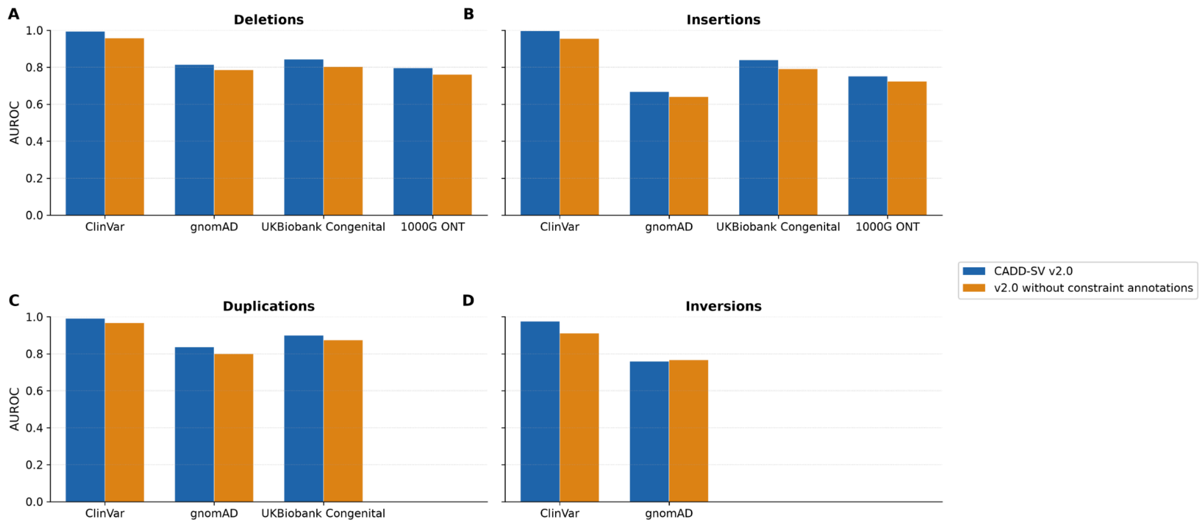


**Supplementary Figure 6: Performance of CADD-SV without evolutionary constraint annotations.** Bars show the AUROC on the ClinVar, gnomAD, UK Biobank congenital-disorder (UKBB) and 1000 Genomes ONT (1000G ONT) validation sets, where available. Deletions ***(A)***, insertions ***(B)***, duplications ***(C)*** and inversions ***(D)*** are evaluated separately. For the model without constraint annotations all features deriving from PhastCons, Zoonomia, GERP, Ultraconserved elements, Microsynteny, LINSIGHT, CADD, MPC, CCR and pLI were removed. Despite removing those features the model still retains good performance.

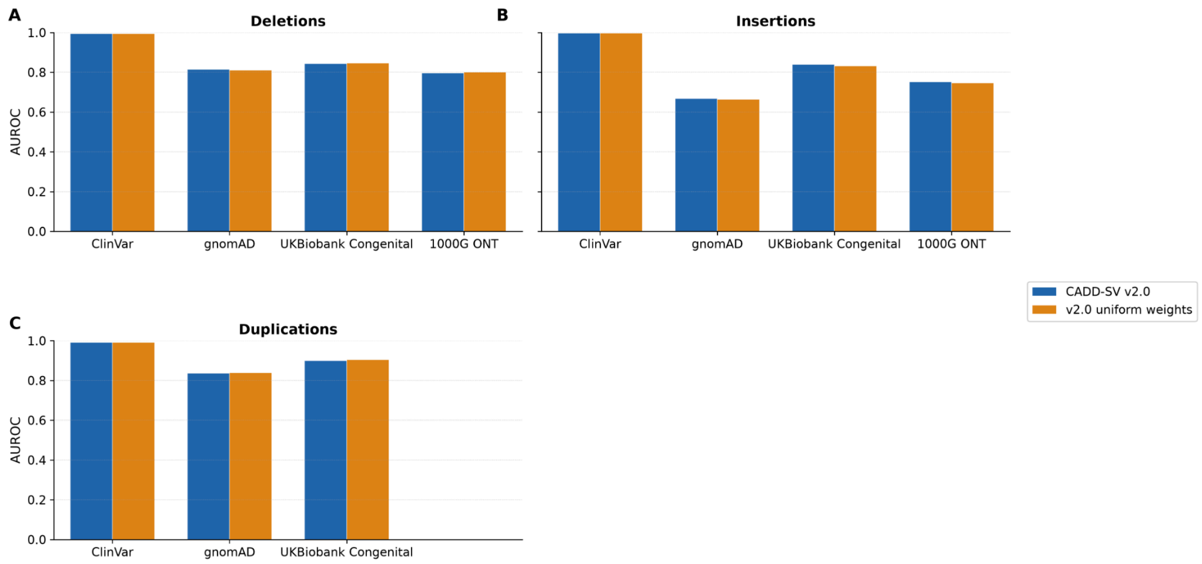


**Supplementary Figure 7: Impact of insertion weighting on CADD-SV v2.0 performance.** Bar plots compare the area under the receiver operating characteristic curve (AUROC) for the standard CADD-SV v2.0 model (blue) and a model trained with uniform sample weights (orange). Results are shown separately for ***(A)*** deletions, ***(B)*** insertions and ***(C)*** duplications across ClinVar, gnomAD, UK Biobank congenital-disorder (UKBB) and 1000 Genomes ONT (1000G ONT) validation datasets, where available. The weighting comparison assesses the effect of addressing the relative underrepresentation of insertions in the training data. Performance is broadly similar between the two weighting configurations across the evaluated variant classes and datasets, with only small differences in AUROC.

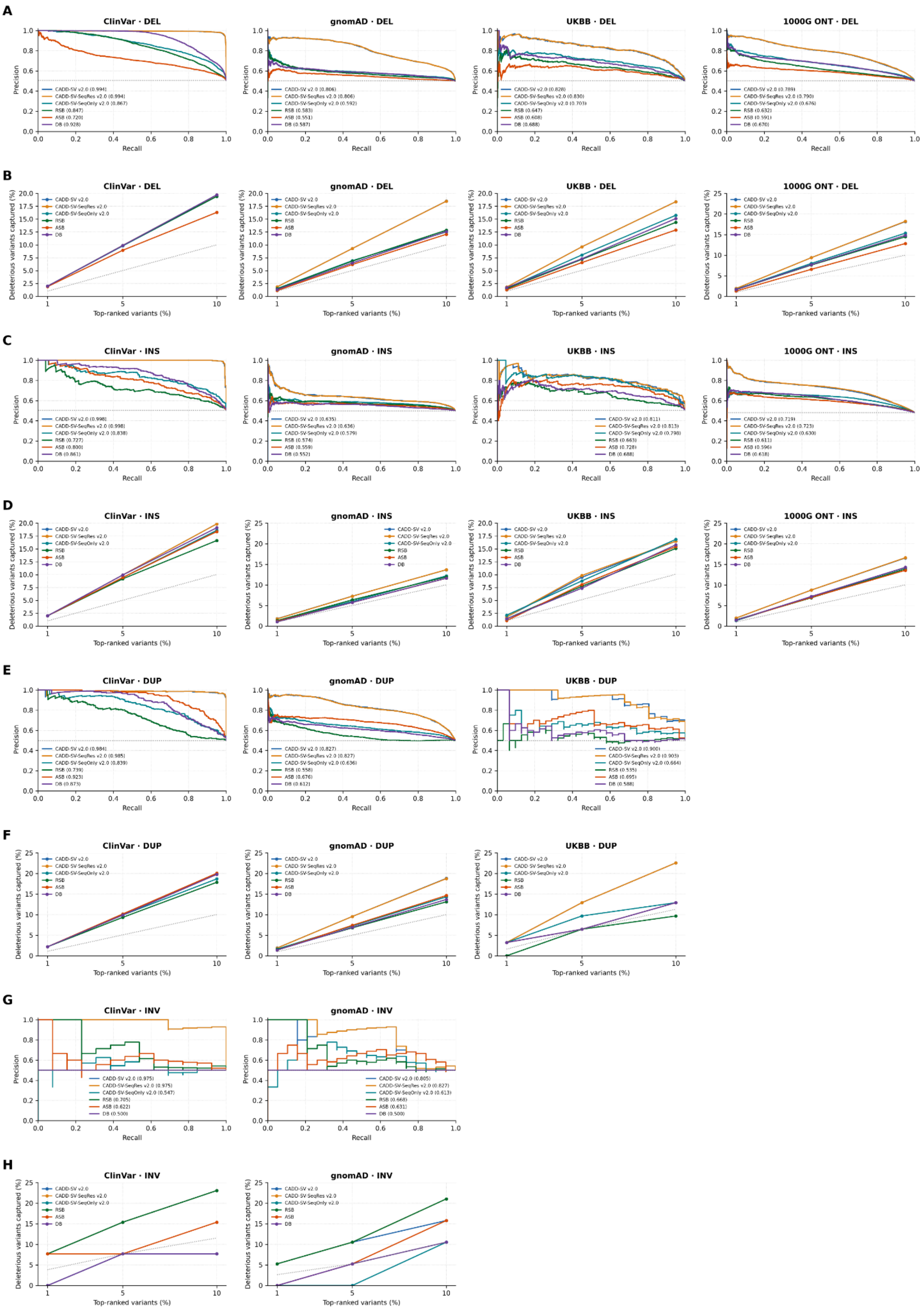


**Supplementary Figure 8: Precision-recall performance and top-ranked variant enrichment across CADD-SV v2.0 model variants and baseline scores.** Precision-recall curves (***A, C, E, G***) and the percentage of deleterious variants recovered among the top-ranked 1%, 5% and 10% of variants (***B, D, F, H***) are shown for deletions (***A, B***), insertions (***C, D***), duplications (***E, F***) and inversions (***G, H***). Columns represent the ClinVar, gnomAD, UK Biobank congenital-disorder (UKBB) and 1000 Genomes ONT (1000G ONT) validation datasets, where available. The three CADD-SV model configurations (CADD-SV v2.0, CADD-SV-SeqRes v2.0 and CADD-SV-SeqOnly v2.0) are compared with the intermediate models (RSB, ASB and DB). Precision-recall legends report average precision, and dotted reference lines indicate baseline expectations. The full and sequence-resolved models show broadly similar performance in many benchmarks, whereas the sequence-only model exhibits greater variation across variant classes and datasets. The relative ranking of models also depends on the benchmark and the fraction of top-ranked variants considered.

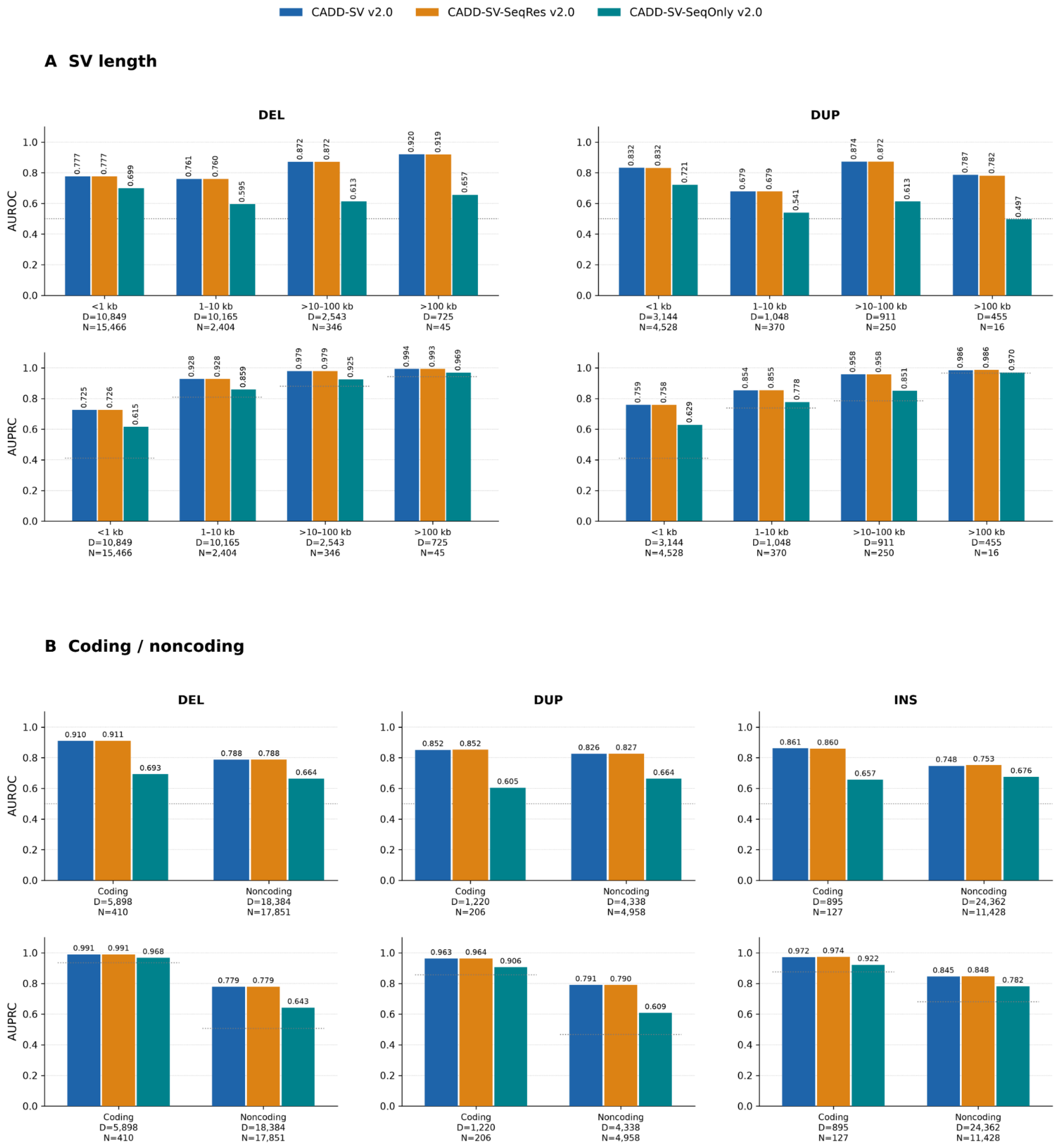


**Supplementary Figure 9: Performance of CADD-SV v2.0 models across structural variant size classes and coding/noncoding regions.** Bar plots compare CADD-SV v2.0 (blue), CADD-SV-SeqRes v2.0 (orange) and CADD-SV-SeqOnly v2.0 (teal) across pooled validation datasets. ***(A)*** SV length: Performance for deletions (DEL) and duplications (DUP), stratified by variant length (<1 kb, 1-10 kb, >10-100 kb and >100 kb). ***(B)*** Coding/noncoding: Performance for DEL, DUP and insertions (INS), stratified by overlap with exons (including UTRs) of protein-coding transcripts using Ensembl release 96 GRCh38 annotations. Variants were classified as coding if their reported reference interval overlapped at least one base of these exons, and noncoding otherwise. Discrimination is measured by the area under the receiver operating characteristic curve (AUROC; upper row of each panel) and the area under the precision-recall curve (AUPRC; lower row). Values above bars show performance estimates, and labels below each category report the number of deleterious (D) and neutral (N) variants. Dotted reference lines indicate AUROC = 0.5 or the stratum-specific deleterious fraction for AUPRC. The standard and sequence-resolved models perform similarly across categories, the sequence-only model generally shows lower discrimination. In panel ***(A)***, AUPRC generally increases with variant length, although larger bins contain fewer examples and should therefore be interpreted cautiously. In panel ***(B)***, performance is generally higher for coding-labelled than noncoding-labelled variants, particularly for AUPRC. Differences across strata should be interpreted alongside variation in class prevalence and validation-dataset composition.

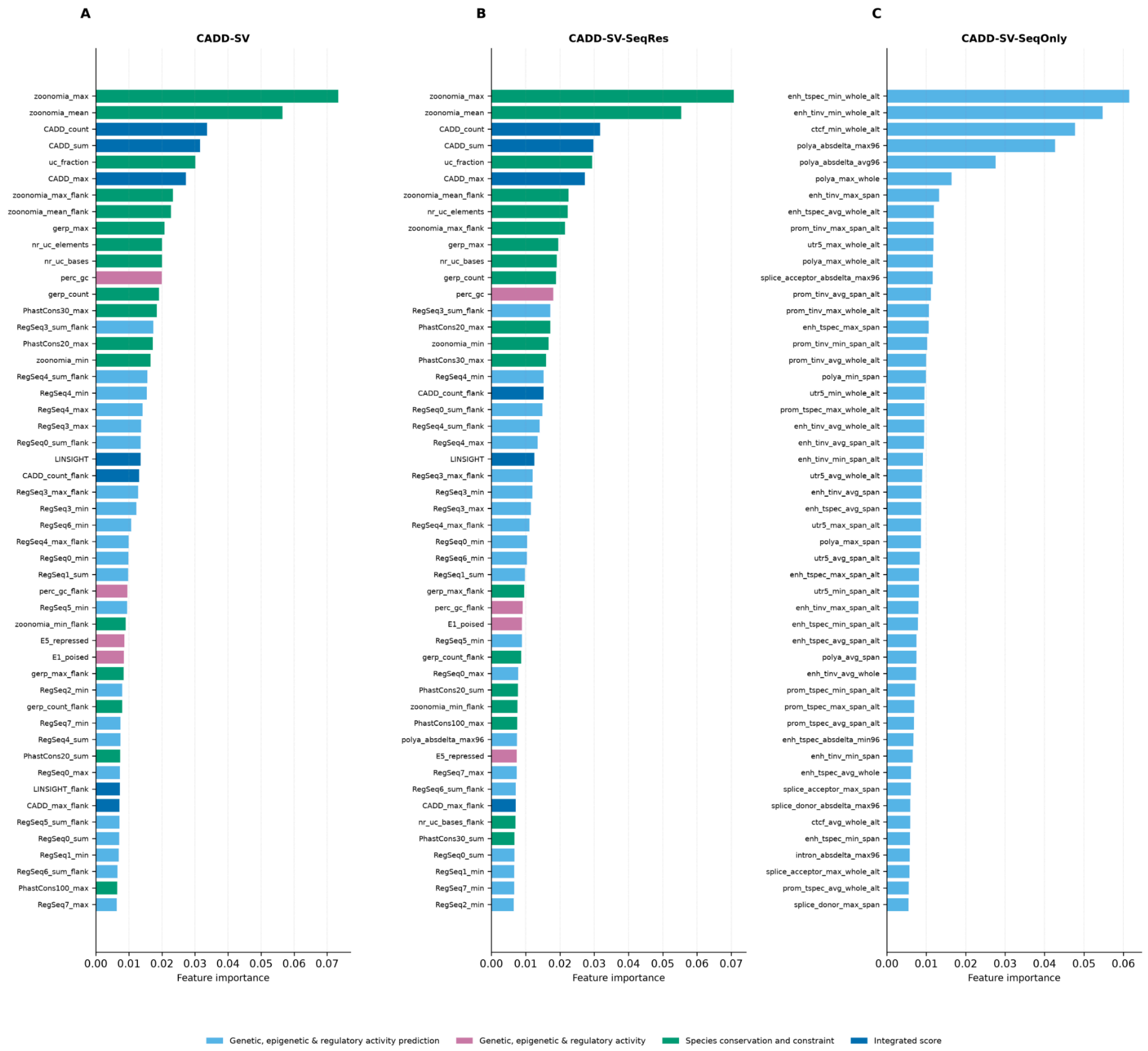


**Supplementary Figure 10: Feature importance profiles across three CADD-SV v2.0 model variants.** Bar plots show the top 50 most important features, ranked by their contribution to model predictions (x-axis: feature importance), for the three models: ***(A)*** CADD-SV v2.0, ***(B)*** CADD-SV-SeqRes v2.0, and ***(C)*** CADD-SV-SeqOnly v2.0. Features are color-coded by category: species conservation and constraint (green), integrated scores (dark blue), genetic, epigenetic and regulatory activity (pink), and genetic, epigenetic and regulatory activity predictions (light blue). Across models incorporating conservation information (A, B), phylogenetic conservation metrics derived from Zoonomia (i.e., zoonomia_max and zoonomia_mean) are consistently the most influential features, indicating that cross-species conservation and constraint provide major predictive signals for structural variant impact. Integrated scores such as CADD_sum and CADD_count, together with additional conservation and constraint metrics (e.g., uc_fraction, gerp_max and nr_uc_elements), also contribute substantially. The CADD-SV-SeqRes model (B) exhibits a feature importance profile broadly similar to that of the original CADD-SV model (A), with conservation features retaining their dominant contribution. Notably, polya_absdelta_max96 is the only sequence-derived prediction feature among the top 50 features in model B, suggesting that sequence-based predictions contribute comparatively little to the overall feature importance in this model. In contrast, the CADD-SV-SeqOnly model (C) relies exclusively on predicted genetic, epigenetic and regulatory activity features. Its highest-ranked features include enhancer activity predictions (e.g., enh_tspec_min_whole_alt and enh_tinv_min_whole_alt), CTCF binding predictions (ctcf_min_whole_alt), and polyadenylation-related predictions (e.g., polya_absdelta_max96 and polya_absdelta_avg96). Additional contributions arise from promoter activity, transcription-associated regulatory predictions, and splice-site signals, highlighting the range of predicted regulatory and transcript-processing features used by the sequence-only model.

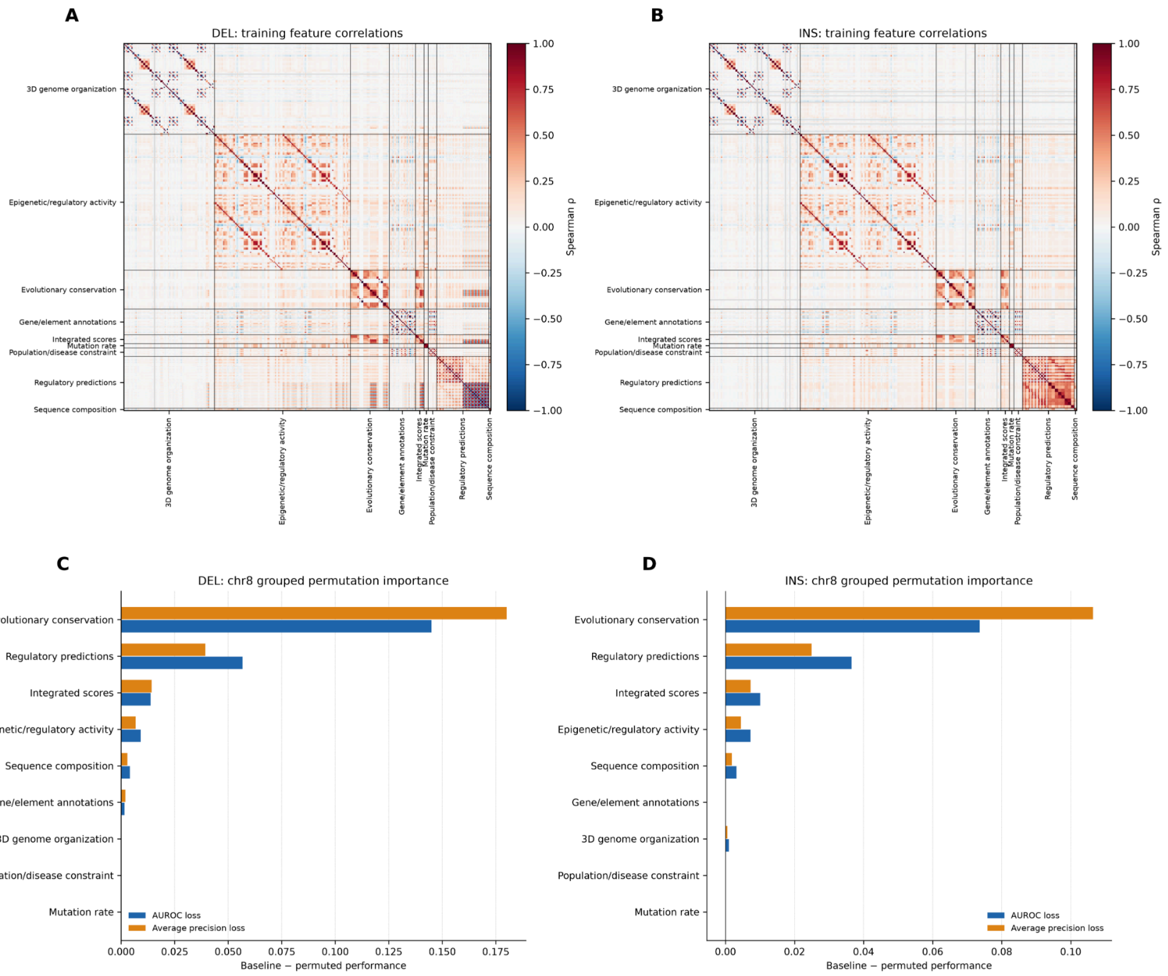


**Supplemental Figure 11: Feature correlations and grouped permutation importance for CADD-SV v2.0 default model.** Spearman correlation matrices for features used to train the deletion ***(A)*** and insertion ***(B)*** models are organized by annotation category, including 3D genome organization, epigenetic/regulatory activity, evolutionary conservation, gene/element annotations, integrated scores, mutation rate, population/disease constraint, regulatory predictions and sequence composition. Red and blue indicate positive and negative correlations, respectively, with the color scale spanning -1 to 1. Grouped permutation importance was evaluated on chromosome 8 for deletions ***(C)*** and insertions ***(D)***. Horizontal bars indicate the loss in AUROC (blue) and average precision (orange) after permuting features within each annotation category; larger decreases indicate greater predictive contribution. Evolutionary conservation produces the largest performance decrease for both variant classes, followed by regulatory predictions, while the remaining categories have smaller effects.

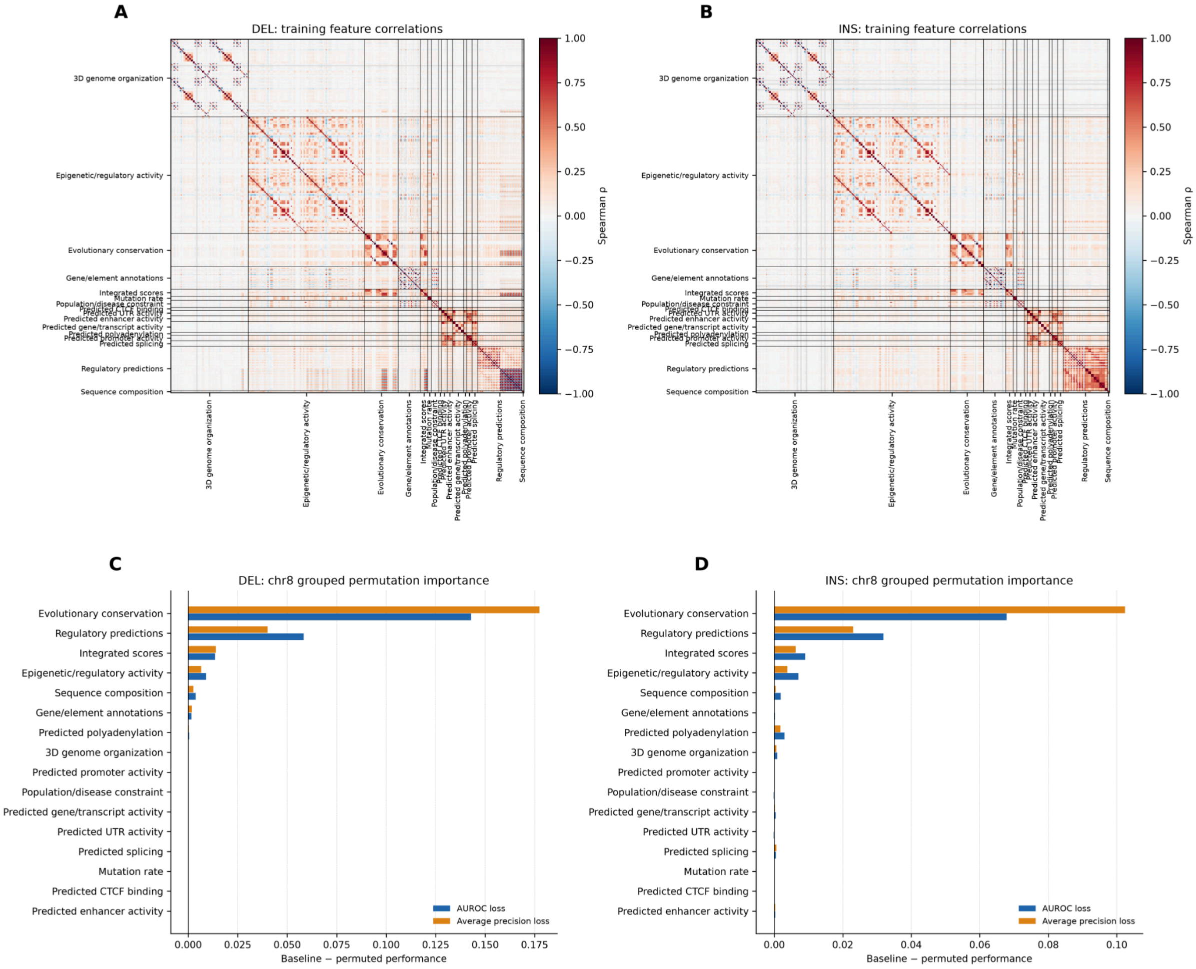


**Supplementary Figure 12: Feature correlations and refined grouped permutation importance for CADD-SV-SeqRes v2.0 model.** Spearman correlation matrices show pairwise relationships among training features for deletion ***(A)*** and insertion ***(B)*** models. Features are grouped by functional annotation category, with sequence-derived predictions further subdivided into categories such as predicted enhancer, promoter, gene/transcript and untranslated-region activity, CTCF binding, polyadenylation and splicing. Red denotes positive correlation and blue negative correlation (Spearman ρ from -1 to 1). Chromosome 8 grouped permutation importance is shown for deletions ***(C)*** and insertions ***(D)***, measured as the decrease in AUROC (blue) or average precision (orange) relative to the unpermuted model. Evolutionary conservation remains the dominant feature group, with regulatory predictions providing the next-largest contribution. Individually separated sequence-derived categories show comparatively small permutation effects in these models.

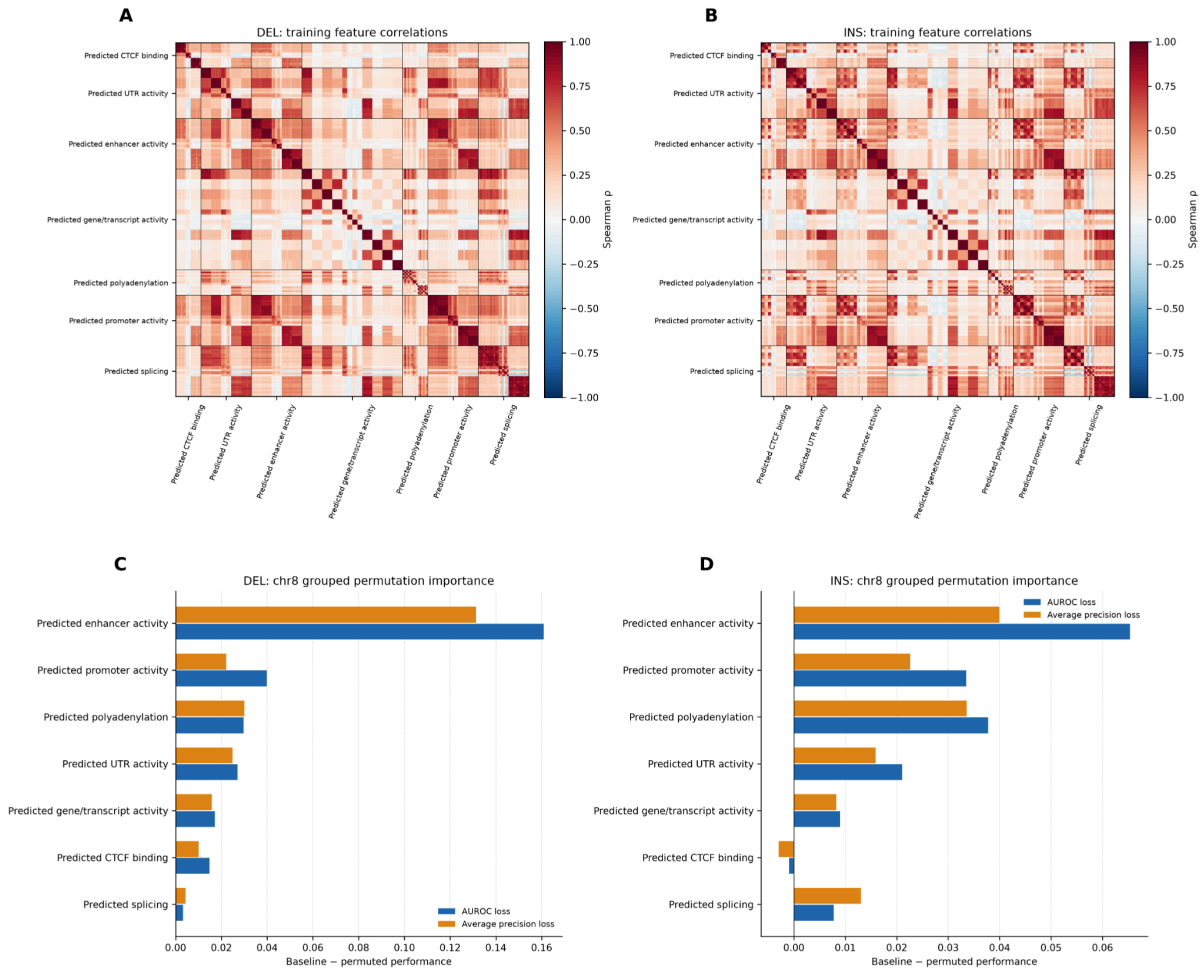


**Supplementary Figure 13: Feature correlations and refined grouped permutation importance for CADD-SV-SeqOnly v2.0 model.** Spearman correlation matrices for predicted regulatory and transcript-processing features are shown for deletion ***(A)*** and insertion ***(B)*** training sets. Feature groups include predicted CTCF binding, untranslated-region (UTR) activity, enhancer activity, gene/transcript activity, polyadenylation, promoter activity and splicing. The color scale represents Spearman correlation coefficients (ρ) from -1 (blue) to 1 (red), revealing blocks of correlated predictions within and across functional groups. Chromosome 8 grouped permutation importance for deletions ***(C)*** and insertions ***(D)*** is quantified by the reduction in AUROC (blue) and average precision (orange) when each feature group is permuted. Predicted enhancer activity is the most influential group for both variant classes; predicted promoter activity, polyadenylation and UTR activity make additional contributions, whereas CTCF-binding and splicing predictions generally show smaller effects.
